# Competing constraints on protein availability and nutrient uptake reshape yeast genetic interactions

**DOI:** 10.64898/2026.09.11.750905

**Authors:** Eivind Almaas, Tjaša Kumelj

## Abstract

Epistasis predicted by enzyme-constrained metabolic models is computed at a single value of the protein available to metabolism, although that amount varies two-to three-fold with growth rate. We ask how epistatic interactions are affected by computing every single and pairwise gene deletion in an enzyme-constrained model of *Saccharomyces cerevisiae* while sweeping protein availability, with and without a bound on nutrient uptake. Protein availability changes relative fitness only under co-limitation, when the protein pool and a nutrient constraint limit growth together. The pool alone, the configuration in which these models are calibrated, makes protein a mere scale factor: no interaction changes sign. Under co-limitation, one genetic interaction in three is gained or lost across the physiological range. Most of this is inherited from single mutants of the respiratory machinery, which is recruited as protein becomes available. The epistasis between a glycolytic lesion with a protein-expensive bypass and the respiratory chain changes sign where the optimum switches from fermentation to respiration. We also find protein-priced interactions: the network supplies a bypass around each deletion and the pool gives it a price, so two genes with no stoichiometric coupling interact whenever the bypass around one competes with the route of the other for the protein pool. Isozyme pairs predicted neutral by flux balance analysis show negative epistasis for the same reason. Relative fitness depends on protein availability and nutrient supply only through their ratio. Thus, nutrient supply is a directly accessible experimental parameter to test epistatic interactions. Sign inversion, isozyme pairs, and interactions with cofactor-consuming branches are predictions a condition-resolved genetic interaction screen can test.

**Author summary:** Enzymes are protein, and a cell has only a limited amount. The share a yeast cell can spend on metabolic enzymes varies two-to three-fold between slow and fast growth. Metabolic models include this limit, yet usually hold it at one fitted value when investigating genetic interactions: whether deleting two genes together is more or less harmful than expected from the separate deletions.

We recomputed every single and double deletion while varying the protein available to the modeled yeast metabolism. Changing protein availability increased growth in mutants and wild type almost equally when protein was the only limitation and left their interactions unchanged. About one-third of interactions appeared or disappeared across the physiological range once glucose uptake also limited growth. Additionally, the model predicts that one class of interaction reverses sign when the cell shifts from fermentation to respiration. Other interactions linked genes in different pathways: when the network offers a detour around each deletion, the enzyme limit gives every detour a cost, and a detour paid with enzyme that another reaction needs makes the double deletion cost more, or less, than both singles together. This simulation-based prediction could be tested experimentally by comparing genetic interactions under controlled high and low glucose supply.

## Introduction

Constraint-based models of genome-scale metabolic networks predict phenotypes from stoichiometry and a small set of inequality constraints [1, 2]; the constraints imposed determine which phenotypes a model can express. Flux balance analysis (FBA) bounds nutrient uptake, whereas enzyme-constrained formulations add a bound on the total mass of catalytic protein (FBA with molecular crowding [3, 4], MOMENT [5] and its simplified form sMOMENT [6], CAFBA [7] and GECKO [8, 9]), coupling each reaction flux to the abundance and turnover number of the enzyme that carries it. This addition reproduces phenomena that FBA cannot, most prominently overflow metabolism, and it has been widely adopted [9–11]. Calibrating such a model sets one additional quantity: protein availability. The procedure allows the saturation factor to be refitted for each condition studied [8], but in practice, this factor is determined once and held fixed across every condition a study explores.

Protein availability varies across physiological conditions. In aerobic, glucose-limited chemostats, the protein content of *S. cerevisiae* biomass rises with the dilution rate, from 38% of dry weight at *D* = 0.025 h^−1^ to 53% at the onset of fermentation and 46% above it [12], and within that protein the fraction allocated to glycolytic enzymes and chaperones falls linearly with growth rate as translation takes a larger share [13]. Rich media reallocate proteome from amino acid biosynthesis to ribosomes [14], nitrogen limitation exposes large unused metabolic capacity [15], and the environmental stress response redirects expression away from growth-related functions [16, 17]. Together, these observations imply substantial, potentially two-to three-fold variation in metabolic protein availability within a single organism’s physiological range, even before heterologous-expression burden or differences between strains are considered.

How this range affects a cell’s genetic interactions is unknown. Genetic interactions, the dependence of one mutation’s fitness effect on the presence of another, are the standard probe of functional organization in metabolic networks [18, 19]. Theory for how epistasis propagates through such a network predicts that it should depend on the genetic background and on the environment [20], and screens report both, for the phenotype scored [21] and, for a minority of pairs, for the growth condition [22]. Interactions are nonetheless computed almost exclusively at a single, fixed protein availability. Whether that choice is innocuous is a question about the structure of the models themselves. Under the assumptions of elementary-flux-mode analyses, growth-rate maximization with a single growth-limiting constraint selects an optimal extreme mode; a gradual shift between distinct metabolic strategies therefore requires at least two constraints to bind simultaneously [23–25]. With the protein pool as that single constraint, the predicted growth rates are linear in protein availability [10]. The configuration in which enzyme-constrained models are calibrated is exactly this one, the saturation factor being fitted to a measured maximum growth rate with nutrient uptake unconstrained [8].

There is also reason to expect that any resulting interactions will differ from those predicted by FBA. In a stoichiometric model, two genes interact through flux coupling: their reactions share a metabolite, or steady-state mass balance couples them through intervening reactions [26]. This coupling follows the connectivity of the network, which helps explain why FBA interactions are modular and concentrated within and between pathways [18]. An enzyme constraint adds a coupling of a different reach. It gives every link in the network a cost, the enzyme mass that a unit of flux requires, and it fixes the total. Flux rerouted onto one link is therefore enzyme taken from other links, wherever the pool binds. A deletion then costs what the cheapest remaining route costs. Two deletions interact when both of their routes draw on the pool at once, or when one deletion removes the route its partner competes with. The topology of the network decides which pairs have such routes, and the pool decides what the routes cost. Interactions generated by this global coupling should appear between genes without direct stoichiometric coupling and should be absent from FBA, which assigns no cost to enzyme use.

This raises the interesting question: under what conditions can protein availability change a genetic interaction at all, and when it can, what does it change? We address it for *S. cerevisiae* metabolism with an enzyme-constrained genome-scale model, sweeping protein availability over a wide range of values, including those a cell visits, and computing every single and pairwise deletion at each value.

We show that the answer has three parts. (1) Protein availability changes relative fitness only under *co-limitation*, when the protein pool and a bound on nutrient uptake limit growth together. It changes nothing but a maintenance correction where the pool binds alone, the configuration in which these models are calibrated. (2) Under co-limitation, most of the reorganization is inherited from single mutants of the respiratory machinery, recruited as protein becomes available. (3) The rest is genuinely pairwise: genes with no direct stoichiometric coupling interact because they compete for the pool, an interaction we call *protein-priced*. The perturbation studied here has a direct experimental counterpart in the supply of the limiting nutrient, because protein availability and nutrient supply enter the model only through their ratio.

## Materials and methods

### Enzyme-constrained model and constraint configurations

Flux balance analysis maximizes biomass-reaction flux subject to steady-state mass balance and flux bounds. The GECKO formulation additionally couples each enzyme-catalyzed flux to enzyme abundance and bounds the total enzyme mass [8]. Writing for *N* the set of reactions, ℳ for the metabolites and *G* for the enzymes with assigned turnover numbers, the optimization problem solved throughout is

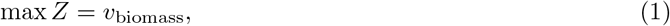

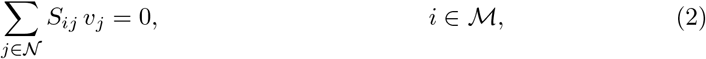

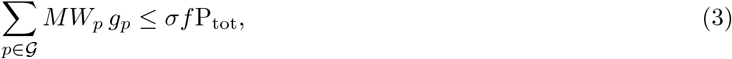

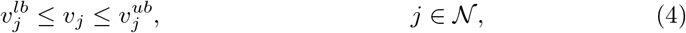

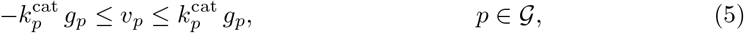

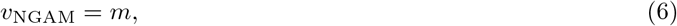

We refer to the total catalytic protein mass on the left-hand side of Eq (3) the *protein pool*, and Eq (3) itself the pool constraint; P_tot_ is the total protein availability that bounds it. We write *ν* for a specific rate in mmol gDW^−1^ h^−1^, the quantity written *q* in the experimental literature, and reserve *v* for the flux variables of the linear program. Here *S*_*ij*_ is the stoichiometric matrix, *g*_*p*_ the concentration and *MW*_*p*_ the molecular weight of enzyme 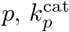 its turnover number, *σ* the average saturation, *f* the fraction of the proteome that is enzyme, P_tot_ the total protein content per gram dry weight, and *m* the non-growth-associated maintenance demand (Eq (6) is a fixed bound in Eq (4), written separately because it plays a distinct role below). We write *µ* for the optimal value of *Z*, the specific growth rate; where its dependence on the parameters matters, it is written out, as in *µ*(P_tot_). Fluxes *v*_*j*_ are in mmol gDW^−1^ h^−1^, *g*_*p*_ in mmol gDW^−1^, *MW*_*p*_ in g mmol^−1^ and 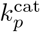 in h^−1^, so both sides of Eq (3) are masses in g gDW^−1^. Note that Eq (5) is written for the one-reaction case. When one enzyme catalyzes several reactions, Eq (5) holds per enzyme–reaction pair, with 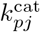 the turnover number of enzyme *p* in reaction *j*. The glucose and oxygen uptake bounds, 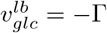 and 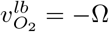, are the only other bounds we vary.

Eqs (2), (3) and (5) are homogeneous of degree one in (*v, g*). If they were the only active constraints, the optimal value would scale exactly with P_tot_; under the relevant assumptions, an optimal extreme-point solution can be represented by an elementary flux mode [23, 27, 28],

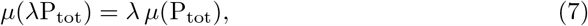

so that every ratio of growth rates, including the relative fitness of every mutant, would be independent of P_tot_ [10]. The maintenance demand of Eq (6) is the one always-active, inhomogeneous constraint. Within each interval with a constant active set, it makes the optimum affine rather than linear,

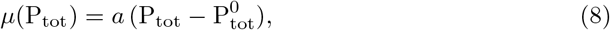

with 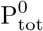 the protein availability consumed in sustaining *m*.

We write *µ*′ for the marginal value of protein: the derivative of the optimum, the slope *a* of Eq (8), and *σf* times the dual variable *y* of the pool constraint, Eq (3),

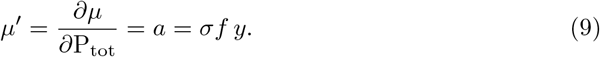

It is constant on each interval of constant active set and steps at the boundaries between them, and it is zero wherever the pool is slack, where added protein buys no growth. At a locally differentiable optimum where the relevant catalytic constraint is active, 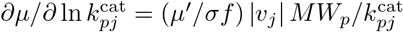 (ecYeastGEM represents each reversible enzyme-catalyzed reaction as two irreversible copies, so every *v*_*j*_ in the model is non-negative and the absolute value is a formality). Thus, the protein mass committed to an enzyme is proportional to the sensitivity of growth to its turnover number, with proportionality determined by *µ*′*/σf*; this is the enzyme-sensitivity relation of protein-allocation models [29, 30]. Eqs (2), (3), (5) and (6) are also jointly homogeneous in (P_tot_, Γ, Ω, *m*): scaling all four by a common factor *λ* scales every optimal flux and the growth rate by *λ*,

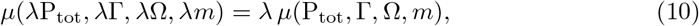

provided every other finite bound is slack as it is at the optima of our configurations. Any ratio of growth rates between genotypes therefore depends on these four quantities only through the ratios Γ*/*P_tot_, Ω*/*P_tot_ and *m/*P_tot_. The consequences for mutant fitness and for the interaction landscape are stated where those quantities are defined. We name each configuration by the constraints that bind at the optimum. The protein pool alone with both uptakes unbounded, as in the GECKO calibration [8]; a nutrient bound alone, as in standard FBA; and the protein pool together with a nutrient bound.

Below the P_tot_ at which the pool constraint starts to bind, the optimum follows the respiratory mode, and every enzyme demand in that solution is proportional to the glucose uptake. The enzyme needed to respire glucose at uptake flux *ν*_glc_ is therefore *c ν*_glc_, where *c* is the protein cost of the respiratory mode per unit glucose flux. The pool becomes limiting where 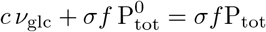, hence

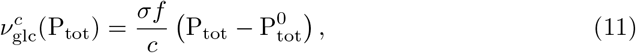

which is linear in P_tot_, with an intercept set by maintenance. We estimate the constant *c* from the model.

### Model adjustment and solver settings

We used the enzyme-constrained model ecYeastGEM v8.6.2 and its parent model, Yeast-GEM v8.6.2 [9, 31]. We restricted the gene set to the 1,159 genes with systematic ORF identifiers (`Y[A-P][LR][0-9][0-9][0-9][WC]`, with or without the `-A` suffix, plus the mitochondrial `Q`-series). Turnover numbers were taken from ecYeastGEM. GECKO assigns them from BRENDA, with cross-organism fuzzy matching where no *S. cerevisiae* value is available [9, 32].

The model implements the pool constraint as the lower bound of a pool-exchange reaction. This bound is the enzyme mass *σf* P_tot_ from Eq (3), expressed in mg per gDW: with *σf* = 0.25 it equals 250 P_tot_, and the bound of the released ecYeastGEM, 125 mg gDW^−1^, corresponds to P_tot_ = 0.5 g gDW^−1^. We therefore report P_tot_ throughout. The physiological range of enzyme mass *σf* P_tot_ is roughly 0.1 to 0.3 g gDW^−1^, i.e. P_tot_ ≈ 0.4 to 1.2. Under the uptake bounds used here, the pool goes slack at P_tot_ = 0.658, and above that value the wild type no longer responds to P_tot_; the physiological window analyzed below, P_tot_ = 0.4 to 0.66, is the part of the physiological range in which the pool still binds.

Unless stated otherwise, we set the uptake bounds to Γ = 5 and Ω = 8 mmol gDW^−1^ h^−1^, and the maintenance demand *m* is kept at its published value, 0.7 mmol ATP gDW^−1^ h^−1^. We closed three exchange reactions in both models. We refer to the model ecYeastGEM v8.6.2 as released as *ecYeastGEM*, and to the same model with these three exchanges closed as *ecYeastGEM-A*. Acetaldehyde (r 1631) is toxic to *S. cerevisiae*, and its efflux is slow enough for intracellular concentrations to exceed extracellular concentrations [33, 34]. (R,R)-2,3-butanediol (r 1549) is a minor fermentation product, formed from acetoin by the dehydrogenase Bdh1p [35, 36], that reaches appreciable titres only in strains engineered for it [37]. Diphosphate (r 4527) is a by-product of biosynthetic reactions whose hydrolysis by a soluble cytosolic pyrophosphatase keeps those reactions running forward; it is not excreted [38]. Results describes the consequences of leaving these routes open.

All simulations used reframed 1.5.3 with Gurobi 10, with FeasibilityTol 10^−8^, OptimalityTol 10^−9^ and NumericFocus 3. These tight tolerances were required because, under the default settings, deletions of genes whose reactions carry ~ 10^−7^ mmol gDW^−1^ h^−1^ of essential flux (heme, thiamine, the GABA shunt) are mis-scored as neutral, which affected multiple genes in a preliminary run.

### Sampling P_tot_ values and locating phase boundaries

We solved the ecYeastGEM-A wild type at 1,002 values of P_tot_ (0.004 to 1.0 in steps of 0.001; the values 1.2, 1.6, 2.0, 3.0 and 4.0 were also included to anchor the saturated regime), recording the reduced costs of the pool, glucose and oxygen bounds. A bound is active where its reduced cost exceeds 10^−9^ in magnitude, and a phase boundary is a value of P_tot_ at which the set of active bounds changes. A bound that is binding but whose reduced cost is zero therefore counts as inactive: relaxing it would not change the optimum.

Along the same sweep, we recorded how the pool is allocated at P_tot_ = 0.004 to 1.0 in steps of 0.002 below 0.1 and 0.01 above: the protein mass 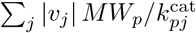 committed to each enzyme *p*, aggregated by the model’s subsystem. By the enzyme-sensitivity relation above, this profile is also the profile of growth sensitivities to the individual turnover numbers. Where the pool constraint is slack, allocation is not unique and is therefore not reported.

We solved single and double deletions in ecYeastGEM-A at 421 values of P_tot_ (0.04 to 0.88 in steps of 0.002), with a wild-type solution at every P_tot_. We excluded genes that were lethal at a given P_tot_ (*µ <* 10^−6^) from the double screen at that value; in practice the same 980 genes were viable at every grid value, so every pair count below refers to the same 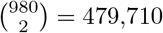 pairs. The screen comprised approximately 4.8 × 10^5^ linear programs per value of P_tot_ and 2.0 × 10^8^ per sweep, totalling 5,800 CPU hours at an average of 103 ms per solution. For the saturation analysis, single deletions were additionally solved from P_tot_ = 0.9 to 2.0 in steps of 0.02 and from 2.1 to 8.0 in steps of 0.1.

For the chemostat scan, we fixed P_tot_, left oxygen unconstrained and varied Γ from 0.25 to 12. The critical uptake 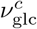 was located by bisection (24 steps) as the smallest bound at which the pool shadow price is non-zero; *µ*^*c*^ is the growth rate at that point. A chemostat at steady state fixes *µ* = *D*, so *µ*^*c*^ is the model’s critical dilution rate, written *D*_crit_. The linear fits of Eq (11) were obtained by ordinary least squares over P_tot_ = 0.30 to 0.70 in steps of 0.05 together with 0.80, 0.90 and 1.00. We located the regime boundaries of the (*ν*_glc_, P_tot_) plane with Ω = 8 mmol gDW^−1^ h^−1^ the same way. At each P_tot_ from 0.04 to 1.00, in steps of 0.005 and 0.001 near the triple point, we found the glucose bound at which the pool, glucose or oxygen constraint switches between slack and binding. The switch was read from the reduced cost and located by bisection on *ν*_glc_ to 5 × 10^−4^ mmol gDW^−1^ h^−1^.

In the comparison with chemostat measurements [12], we estimated P_tot_ in two ways. The first maps the measured *D*_crit_ through the fitted *µ*^*c*^(P_tot_). The second uses least squares against all reported quantities (ethanol, oxygen and carbon dioxide fluxes and biomass yield at ten dilution rates, each normalized by its measured maximum), with model values interpolated at the measured dilution rates. Dilution rates above the model’s maximum growth rate contribute no residual, so the sum runs over 32 to 40 values depending on P_tot_; all 40 are defined from P_tot_ = 0.575 upward, which contains the minimum. These fits reuse one scan: by Eq (10), the scan at any P_tot_ follows from a reference scan by scaling the glucose bound and all fluxes by P_tot_*/*0.596, and the deviations computed this way agree with direct solutions at five values of P_tot_ to within 0.001.

### Quantifying mutant fitness and epistasis as functions of P_**tot**_

All deletion phenotypes were evaluated on the P_tot_ grid defined in the previous subsection, from 0.04 to 0.88 in steps of Δ*P* = 0.002. Extrema over P_tot_ are extrema over this grid. Only the saturation analysis below uses the coarser extension of the grid to P_tot_ = 8. We define the fitness of the deletion of gene *i* as the mutant’s optimal growth rate *µ*_*i*_ normalized to that of the wild type, *µ*_WT_, at the same P_tot_,

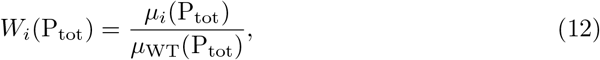

Values were clipped to [0, 1], and values within 10^−4^ of unity were set to unity. Double-deletion fitness *W*_*ij*_(P_tot_) was defined analogously. 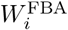 is the fitness in the FBA configuration, the same model and bounds with the pool constraint removed. A single deletion is P_tot_-dependent when its fitness spread over the grid exceeds *K*_*W*_,

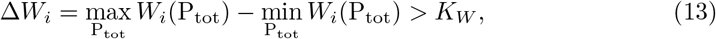

with *K*_*W*_ = 0.05 unless another threshold is stated.

The saturation value 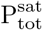 of a strain, either the wild type or the single deletion of gene *i*, is the smallest P_tot_ value in the extended grid at which its growth reaches 99.99% of its value at P_tot_ = 8. That is the same tolerance we use for fitness values at unity. Then

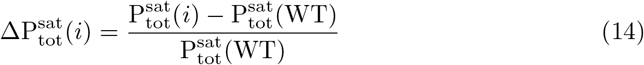

is the additional P_tot_ the deletion strain needs, in units of the wild-type value.

We define the epistatic interaction between genes *i* and *j* as the deviation of the double-deletion fitness from the multiplicative expectation [18],

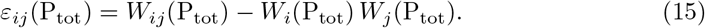

Because Eq (10) scales mutant and wild type alike, fitness obeys

*W*_*i*_(*λ*P_tot_, *λ*Γ, *λ*Ω, *λm*) = *W*_*i*_(P_tot_, Γ, Ω, *m*), and likewise for *W*_*ij*_ and *ε*_*ij*_. Thus, subject to the assumptions underlying Eq (10), the genetic interaction landscape is a function only of the ratios Γ*/*P_tot_, Ω*/*P_tot_ and *m/*P_tot_. We confirmed this numerically, finding that the fitness of each of the 1,159 single deletions at (P_tot_, Γ, Ω, *m*) = (0.25, 2.5, 4, 0.35) and at (1.0, 10, 16, 1.4) agrees with that at (0.5, 5, 8, 0.7) to within 10^−11^. With *m* held at its published value while P_tot_, Γ and Ω are halved or doubled, the largest change in any single-deletion fitness is 0.020 under halving (31 genes above 0.01) and 0.010 under doubling (none above 0.01); this is the maintenance correction to the ratio dependence.

We call a pair interacting at P_tot_ when

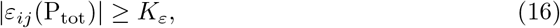

using *K*_*ε*_ = 0.01, with 0.05 as a sensitivity check. The threshold separates a minority of substantial interactions from a mass of values in the range of solver noise: of the 2.02 × 10^8^ pair solutions in the ecYeastGEM-A sweep, 98.99% have |*ε*_*ij*_| *<* 10^−4^, two orders of magnitude below it. We count, classify and compare interactions on the raw *ε*_*ij*_ of Eq (15). For comparison, we also calculate the scaled interaction

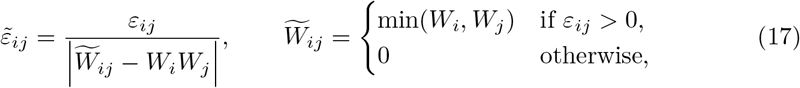

so that 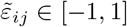. The bounds hold because a second deletion cannot increase the optimum, *W*_*ij*_ ≤ min(*W*_*i*_, *W*_*j*_). It is reported only for interacting pairs. This restriction is numerical: for the great majority of pairs, those below the threshold, the denominator of Eq (17) is itself of the order of solver noise, and 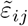 can then take values of order one arbitrarily.

A pair is classified as P_tot_-only if it is interacting at one or more grid values but not in FBA, and persistent if it is interacting at every grid value. A P_tot_-only pair is strict when neither single mutant moves: the fitness of each gene spans less than 0.005 over the grid, half the interaction threshold. Their product *W*_*i*_*W*_*j*_ then cannot vary enough to make the pair interacting on its own. A pair changes sign if it interacts with opposite signs at two grid values; a sign crossing is located at the last grid value at which it interacts with the earlier sign. This was evaluated at *K*_*ε*_ = 0.01 and 0.005. We quantify the local sensitivity of an interaction to P_tot_ by the largest change between adjacent grid values, 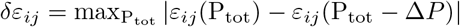. The FBA interaction 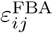 is Eq (15) evaluated on the FBA fitnesses. We recover *µ*′, Eq (9), in every background from the reduced cost of the pool exchange bound returned with the solution: the bound is −250 P_tot_ in mg gDW^−1^, so *µ*′ is 250 times the magnitude of the per-mg reduced cost. We recorded it for every viable deletion at P_tot_ = 0.30, 0.50 and 0.62, and for every pair interacting by Eq (16) at P_tot_ = 0.15, 0.30 and 0.50. For a negative interaction, this gives a first-order price in units of protein availability: the additional protein availability that the double mutant would need, to first order, for its growth to reach the multiplicative expectation is

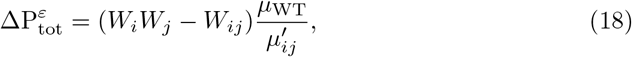

where 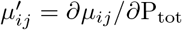 in the double mutant. It diverges as the double approaches the regime where the pool no longer binds, and is undefined beyond it, where no amount of protein closes the deficit. Because dual values need not be unique at a degenerate optimum [39], and because *µ*(P_tot_) is piecewise linear, the single-deletion duals were checked at P_tot_ = 0.5: for all 980 viable deletions, 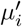 equals a one-sided finite difference of *µ* over ΔP_tot_ = 0.002 to within 10^−11^; the forward and backward differences agree to the same tolerance, so no mutant sits at a kink; and repeating every solution with dual simplex, barrier and default numerical settings changes no 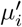 by more than 3 × 10^−9^.

### Assigning genes to pathways within the model

Gene function was taken from the model rather than from external annotation. We assigned each gene the subsystem, among the 106 subsystem groups of ecYeastGEM, that occurs most often among the reactions its gene-protein-reaction rules associate it with. Where that subsystem is a transport group and a metabolic subsystem also occurs, we took the metabolic one. Two genes are model isozymes if they occur in the rule of the same reaction (the enzyme-constrained model expands such reactions into one copy per isozyme, with suffix `EXP n`). We tested each subsystem pair for enrichment among the P_tot_-only interactions with the hypergeometric distribution, drawing from the 479,710 screened pairs. The *p*-values are Bonferroni-corrected over the subsystem pairs observed.

### Testing robustness to turnover-number uncertainty

To test robustness, we multiplied each enzyme coefficient 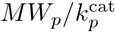 in Eq (5) by 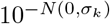, independently per enzyme-reaction pair, for eight replicates each at *σ*_*k*_ = 0.3 and 0.6. We solved single deletions at twenty P_tot_ values per replicate. Saturation thresholds under the same perturbation were located by bisection on the reduced cost of the pool bound, to a tolerance of 10^−6^ in P_tot_.

### Code and data availability

The ecYeastGEM-A files, the deletion-screen tables and the scripts that generate every figure and table in this work are available at https://github.com/AlmaasLab/ecFBA-epistasis; the complete data archive, including the DuckDB databases used to compute the interaction landscapes, is deposited on Zenodo (doi:10.5281/zenodo.22705816). The parent models, ecYeastGEM v8.6.2 and Yeast-GEM v8.6.2, are available from their authors [9, 31].

## Results

Whether protein availability can change a genetic interaction at all is decided by which constraints bind at the optimum (Fig 1). Consider a wild type and a deletion mutant whose only limitation is the amount of metabolically available protein, P_tot_. Doubling that protein doubles every flux in both strains, and each grows twice as fast. The fitness of the mutant relative to the wild type is unchanged, and so is any interaction computed from such ratios (Fig 1A, first two conditions). Now cap the glucose uptake at a rate the wild type already draws, again with double P_tot_. The wild type is now limited by the glucose as well as by the protein: the added protein no longer scales its growth and buys it only a better use of the glucose it imports. The mutant, whose deletion forces it through a more protein-expensive route, stays below the cap and is limited by the protein alone, so it still grows in proportion to it. The wild type therefore gains less than the mutant, and relative fitness increases (Fig 1A, third condition).

**Fig 1.**
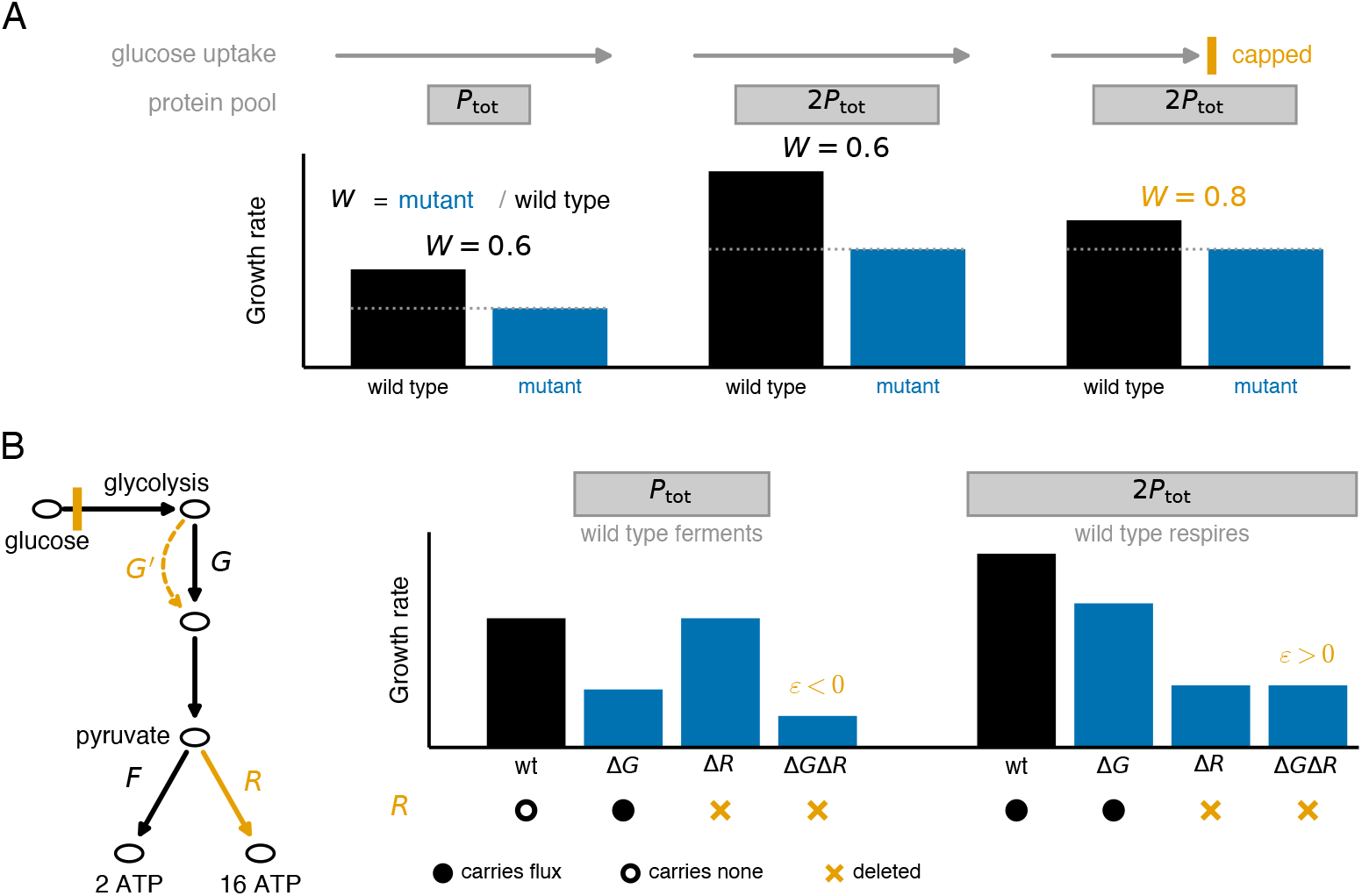
How protein availability can act on genetic interactions under co-limitation. **A**. A mutant against the wild type for three conditions. In the first two, there is only a single bound (protein pool). In the last, there are two bounds (protein pool and glucose uptake). Doubling P_tot_ with glucose uptake unbounded scales both strains and leaves the relative fitness *W* unchanged; doubling P_tot_ with glucose uptake capped at a value the wild type already meets increases *W*. **B**. Left, a toy example for the third condition of **A**: a glycolytic step *G* with a bypass *G*′ (dashed), and pyruvate to ATP by fermentation *F* or respiration *R*. All orange routes are expensive in enzyme. Right, growth of its four genotypes for two values of P_tot_ with the flux state of *R* indicated by symbols below; *F* carries flux in all eight genotypes. *ε* measures the epistatic interaction in the Δ*G*Δ*R* strain.

Applied to a gene pair under that same glucose cap, the reasoning yields a less obvious outcome: it can reverse the sign of an epistatic interaction. Consider a glycolytic step *G* whose deletion is rescued by a bypass *G*′ and a respiratory step *R* that are both expensive in enzyme (Fig 1B). Both draw on the protein pool, so the cost of one depends on whether the other is in use. When protein is scarce (protein pool at P_tot_) the wild type ferments via the step *F*, and reaction *R* carries no flux. Thus, losing *R* costs nothing. However, the Δ*G* mutant must respire with step *R* to make up the ATP that pays for its bypass *G*′. In this situation, losing *R* harms the mutant and not the wild type, and the pair Δ*G*Δ*R* aggravates (Fig 1B, the four genotypes at P_tot_; *ε <* 0).

If instead protein is abundant (protein pool at 2P_tot_), the wild type respires. Deleting *R* now forces fermentation, which is cheap in enzyme, so the protein pool stops binding. The bypass then costs nothing: Δ*G*Δ*R* grows exactly as Δ*R* does, and the pair Δ*G*Δ*R* alleviates (Fig 1B, the four genotypes at 2P_tot_; *ε >* 0). A single parameter, protein availability, thus carries the pair from aggravating to alleviating. Protein availability can then reshape genetic interactions only where a second constraint binds alongside the protein pool, and it is inert where the pool binds alone.

The argument is algebraic. The problem is homogeneous in P_tot_ with the pool as the only binding constraint (Eq (7)). It is also conditional. It holds only while every other bound stays slack and while the maintenance demand stays small beside the growth it supports. Neither is guaranteed once the model leaves the condition at which enzyme-constrained models are calibrated. We therefore test the argument, computing every single and every pairwise deletion over the range of P_tot_, first with the pool acting alone and then alongside a nutrient uptake bound. The second of these demands a model that behaves physiologically far from that calibration condition.

### Closing three secretion routes results in a model that reproduces chemostat physiology

ecYeastGEM is calibrated under one condition: batch growth on excess glucose, with both uptake bounds removed and P_tot_ at its published value of 0.5. Varying P_tot_ = 0.04 to 0.88 with glucose and oxygen uptake constrained at 5 and 8 mmol gDW^−1^ h^−1^ respectively, it produces no ethanol overflow over most of the range, excreting acetaldehyde, (R,R)-2,3-butanediol or diphosphate instead (Fig A in S1 File). None of the three is an overflow product of aerobic, glucose-grown yeast; acetaldehyde, the main one, is undetectable in glucose-limited chemostats below *D*_crit_ [40]. The advantage these exports give the model is 0.32% in growth rate at P_tot_ = 0.5, a near-tie far below the uncertainty of the turnover numbers. Closing the three exchanges (Methods) costs at most 2% of growth, leaves ethanol as the only overflow product across the whole range, and changes nothing at the calibration condition, where neither model uses them. We call the adjusted model ecYeastGEM-A and use it throughout, with the plain-FBA comparison model adjusted the same way.

Protein availability P_tot_ is still a free parameter, and the value carried by the released model belongs to that batch calibration, where the pool acts alone. We need an estimate from a condition in which a nutrient bound competes with the pool. A glucose-limited chemostat is such a condition: the feed sets the glucose uptake and the dilution rate sets the growth rate. Fitting P_tot_ by least squares to the biomass yield, gas exchange and ethanol production that van Hoek and co-workers [12] report at ten dilution rates gives 0.62 g gDW^−1^ (Methods). At that value ecYeastGEM-A reproduces the measurements across the whole dilution-rate range with nothing further fitted (Fig B and Table A in S1 File): the gas fluxes of the respiratory branch to within 0.6 mmol gDW^−1^ h^−1^ and the yield to within 0.03 g g^−1^, the onset of ethanol within one sampling interval of the measured *D*_crit_, and the collapse of the yield above it, with an overshoot of ethanol at the highest dilution rate. Two independent estimates agree with the fit: P_tot_ = 0.60 from the measured *D*_crit_ alone, and 0.62 from the *D*_crit_ of the CEN.PK background on which the model’s biomass composition rests [41]. The measured oxygen uptake falls above *D*_crit_, which van Hoek and co-workers attributed to glucose repression of respiratory enzyme synthesis [12]. The model reproduces the fall without any regulatory term: past the critical uptake the optimum reallocates protein from respiration to fermentation, which yields more growth per unit of enzyme.

### The (*ν*_glc_, P_tot_) plane contains five regimes, and P_tot_ acts on relative phenotypes in only two

We use the ecYeastGEM-A model, first with the pool as the sole capacity constraint and both uptakes unlimited. Homogeneity (Eq (7)) predicts that no relative phenotype depends on P_tot_; only the maintenance demand can spoil this by making growth affine rather than linear (Eq (8)). Over a 250-fold range of P_tot_, we find that growth is affine to numerical precision, *µ* = 0.8269 P_tot_ − 0.0013, and the intercept costs 0.3% of a typical P_tot_.

To identify where a second constraint becomes active, we scanned the glucose bound at P_tot_ = 0.5 with oxygen unlimited, the configuration of an aerobic glucose-limited chemostat (blue dashed line, Fig 2A and Fig 2B). Below 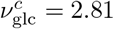 mmol gDW^−1^ h^−1^ the pool shadow price is zero and the enzyme-constrained model returns the growth rate of FBA to four decimal places. Above it, the two curves separate (Fig 2B): the pool binds and ethanol appears (Fig 2A, oxygen stays slack along this line), while FBA overflows at no uptake at all. Repeating the scan across P_tot_ gives 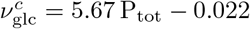 and *µ*^*c*^ = 0.478 P_tot_ − 0.005, both with *R*^2^ = 1.000 (Fig 2A). Whereas the linear form is expected [10, 24], the genome-scale model supplies the constant. Since the enzyme mass behind P_tot_ is *σf* P_tot_ = 0.25 P_tot_, the respiratory mode of ecYeastGEM-A costs 44 mg of enzyme per mmol gDW^−1^ h^−1^ of glucose it oxidizes.

**Fig 2.**
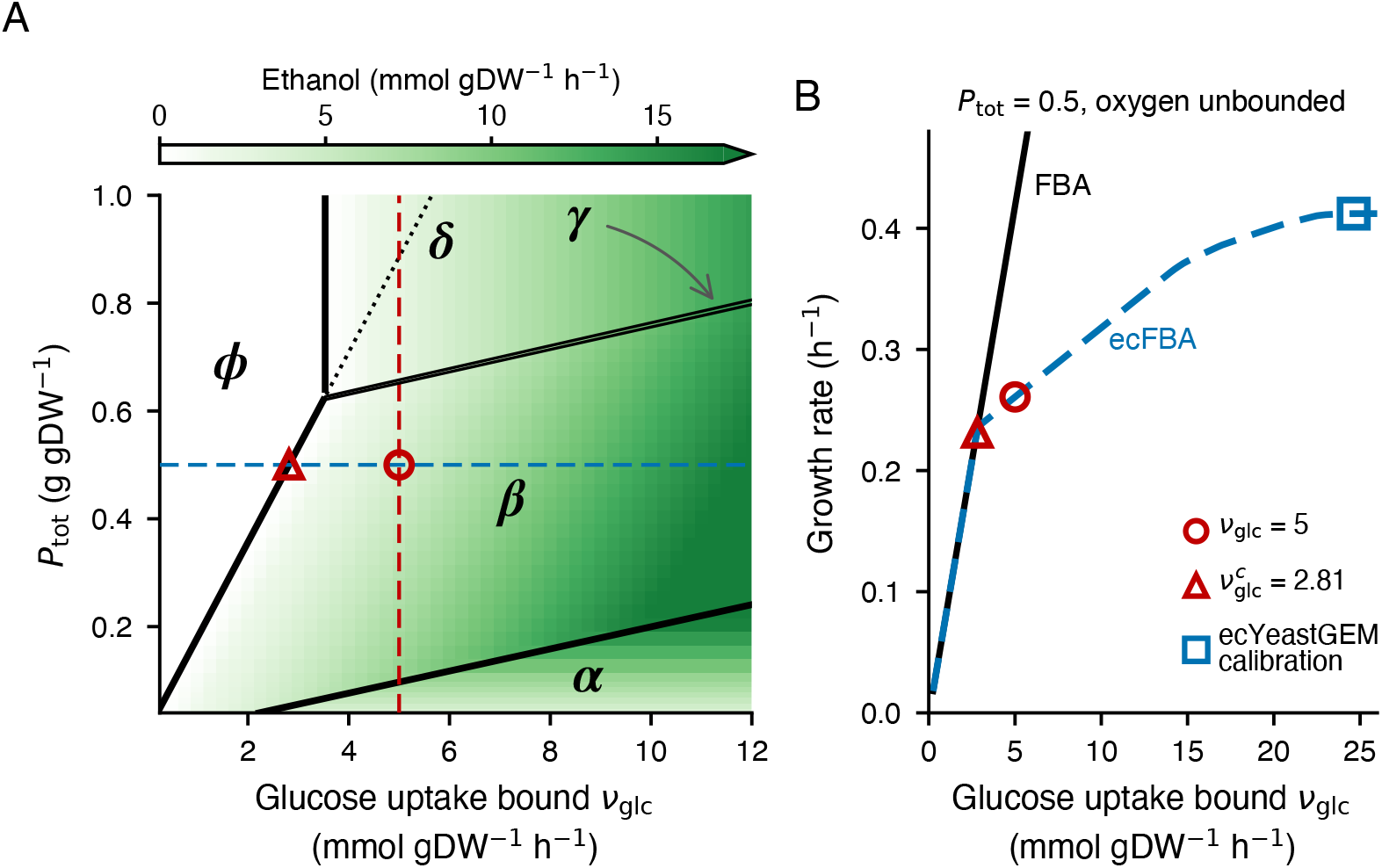
Active-constraint regimes in glucose uptake and protein availability. **A**. Ethanol excretion by the wild type, oxygen bound 8 mmol gDW^−1^ h^−1^. Black lines separate the regimes: *α*, the protein pool binds; *β*, pool and glucose binds; *γ*, all three; *δ*, glucose and oxygen binds; *ϕ*, glucose only. Dotted, 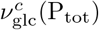 of Eq (11); dashed blue, P_tot_ = 0.5; dashed red, *ν*_glc_ = 5; circle, *ν*_glc_ = 5, and triangle, 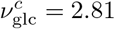, both at P_tot_ = 0.5. **B**. Growth rate against glucose uptake at P_tot_ = 0.5, oxygen unlimited: FBA prediction, black; ecYeastGEM-A, dashed blue; circle and triangle as in **A**; square, the ecYeastGEM calibration with both uptakes unbounded.

The oxygen bound adds a third constraint, and with it the (*ν*_glc_, P_tot_) plane separates into five regimes according to which combination of constraints bind. The pool binds alone in *α*, together with the glucose bound in *β*, and with both nutrient bounds in the narrow wedge *γ* (Fig 2A). The glucose and oxygen bounds are active in *δ*, and the glucose bound alone in *ϕ*. In the phases *ϕ* and *δ*, the pool is slack and the wild type solves the same problem as FBA, to numerical precision in growth rate and every exchange flux. What separates the two is the oxygen bound, which allows complete respiration only up to *ν*_glc_ = 3.53 mmol gDW^−1^ h^−1^. The ethanol of *δ* is therefore oxygen overflow, and FBA excretes exactly the same. The ethanol of *β* is produced by the pool constraint, at oxygen uptakes far below the bound. Relative phenotypes thus depend on P_tot_ only in *β* and *γ*, where the pool is active together with a nutrient uptake bound. In *α*, everything scales with P_tot_ by homogeneity, and in *ϕ* and *δ* P_tot_ does nothing at all. We refer to *β* and *γ* as *co-limitation*, and it is the only part of the plane in which the questions of this paper have a non-trivial answer (Fig 1).

The regimes of Fig 2A are the wild type’s, and every genotype has its own such phase plane: a deletion potentially changes the enzyme needed to carry a given flux, so its boundaries may lie elsewhere in the same plane, and a double or triple deletion has boundaries of its own again. Two strains at one point in the plane can therefore differ in which constraints are active, and a mutant can end with an active set that the wild type never has anywhere along the path we sweep: not a different position on this map, but a different map. As in a traditional phenotype phase plane [42], the regimes are defined by which shadow prices are non-zero, so the duals returned with any single solution classify that genotype at that point without a map of its own.

Varying nutrient supply at fixed P_tot_ (blue-dashed line, Fig 2A) and varying P_tot_ at fixed supply (red-dashed line, Fig 2A) trace orthogonal paths through this plane. Eq (10) makes them equivalent after rescaling, however. Every mutant fitness and every interaction depends on P_tot_ and the uptake bounds only through their ratios, up to a maintenance term worth at most 0.02 in fitness over a two-fold change (see Methods). Up to the maintenance term *m*, our P_tot_ sweep is equivalent to a sweep of nutrient supply at a fixed proteome: halving P_tot_ at *ν*_glc_ = 5 has the same effect as doubling the glucose supply at P_tot_ = 0.5.

We vary P_tot_ at *ν*_glc_ = 5, the respirofermentative regime observed above *D*_crit_ [12]. The sweep traverses *α, β, γ* and *δ* and never enters *ϕ* (red dashed, Fig 2A), and the active set changes at P_tot_ = 0.097, 0.652 and 0.658 (Fig C in S1 File). Through *β* the marginal value of protein falls in steps, each a change in the set of marginal enzymes, as respiration is recruited in stages. The first stage does not make ATP: it replaces the glycerol sink for surplus cytosolic NADH with electron transport, the same sink an evolved Crabtree-negative strain sheds by a truncating mutation in GPD1 [43]. The order of the later stages is not determined, since perturbing the turnover numbers (Methods) shifts them by more than 0.1; what is robust is the composition of the pool, in which the share of oxidative phosphorylation rises from under 1% to a third across *β* in every turnover-number replicate. In *δ* the wild type is identical to FBA, but a deletion strain needs more enzyme to carry the same flux, and just inside *δ*, 34 of the 980 viable deletions still differ from their FBA fitness by more than 0.01.

The largest fitness spread over the whole range with the pool acting alone is Δ*W*_*i*_ = 0.052 (Table 1). The deletions that depend on P_tot_ at all are enzymes of lower glycolysis and the hexose transporter HXT5, which supply maintenance ATP; its cost is fixed, so it is diluted as P_tot_ rises. The dependence is no longer monotone under a glucose bound: for one deletion in six, *W*_*i*_ reverses direction within the P_tot_ range because the constraints that bind change. A pair interacting at one P_tot_ need not interact at another, and not one pair changes sign with the pool alone, against 596 under the glucose bound. In the following, we keep both constraints active.

**Table 1.** Consequence of the P_tot_ constraint on single and double mutants Fitness spread Δ*W*_*i*_, Eq (13), with the threshold *K*_*W*_ varied over the first three rows; pairs interacting by Eq (16). The two columns sample P_tot_ differently: at 0.1, 0.5 and 2.0 with both uptakes unbounded (pool only), and over [0.04, 0.88] with glucose bounded at 5 mmol gDW^−1^ h^−1^.

|  | protein pool only | pool + glucose |
| --- | --- | --- |
| *Single deletions: # of genes with $\Delta W_i > K_W$* |  |  |
| $K_W = 0.01$ | 9 | 85 |
| $K_W = 0.05$ | 3 | 57 |
| $K_W = 0.30$ | 0 | 39 |
| largest $\Delta W_i$ | 0.052 | 0.739 |
| monotone in $P_{\text{tot}}$ | 100% | 83.5% |
| *Interacting pairs* |  |  |
| interacting at $P_{\text{tot}} = 0.5$ | 632 | 1,971 |
| interacting at every $P_{\text{tot}}$ sampled | 628 of 634 | 327 of 2,981 |
| changing sign along the sweep | 0 | 596 |

### P_tot_-dependent deletion phenotypes are the recruited machinery

If P_tot_ acts by recruiting respiratory capacity, the deletion phenotypes that depend on it should be those of the machinery recruited. Of the 1,159 single deletions, 57 are P_tot_-dependent by Eq (13) (Fig 3A), and 47 of them fall into four groups by the model’s subsystem assignment: the subunits of complexes III and IV, those of ATP synthase, the enzymes of lower glycolysis, and pyruvate dehydrogenase, succinate dehydrogenase and fumarase. In *α* every curve is flat apart from the nine maintenance-ATP suppliers of Table 1. Note that the phase boundaries drawn in Fig 3 belong to the wild type. They are relevant to every curve since the strain fitness *W*_*i*_ (Eq (12)) and epistasis *ε*_*ij*_ (Eq (15)) depend on wild-type properties. However, these lines are not the phase boundaries of the strains: each single and each double deletion has its own active set that changes at different values of P_tot_, and most of the bends in Fig 3A fall between the drawn lines rather than on them. Which constraints are active for a mutant therefore has to be read from that strain’s own duals, as below, and not from the wild type’s phase plot (Fig 2A). The set is robust to the turnover number values: perturbing every *k*^cat^ by a lognormal factor with *σ*_*k*_ = 0.3 retains 51 of the 57 on average over eight replicates, and *σ*_*k*_ = 0.6 retains 41 (Methods).

**Fig 3.**
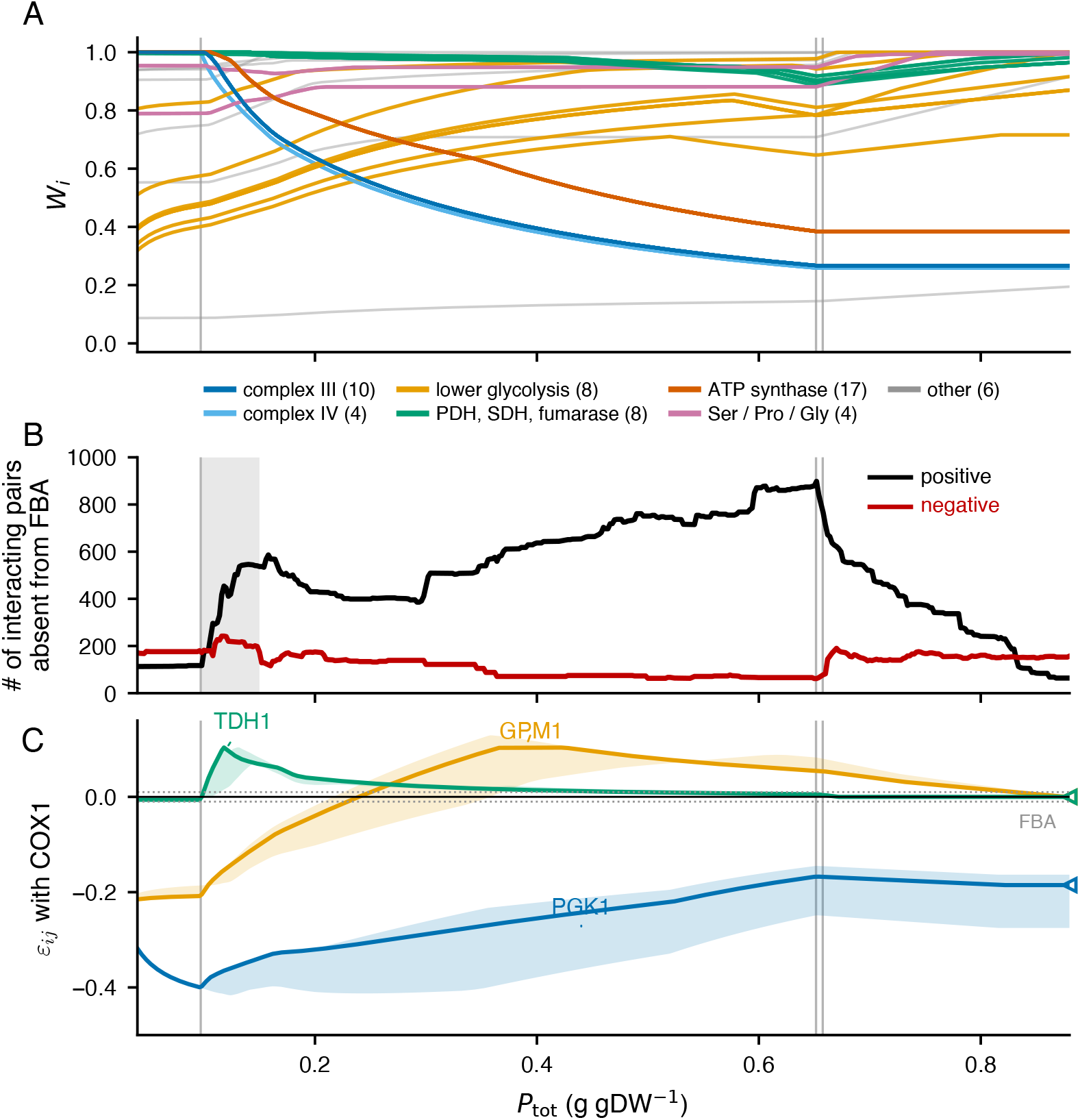
Deletion phenotypes and their interactions along the P_tot_ sweep. (gray lines indicate the wild-type phase-boundary crossings). **A**. *W*_*i*_ for the 57 P_tot_-dependent single deletions, colored by subsystem with the number of genes in the key; the six outside the named groups are pale. **B**. Positive (black) and negative (red) interacting pairs at each P_tot_ with |*ε*_*ij*_| ≥ 0.01, counting only pairs that never interact in FBA; shaded, P_tot_ = 0.097 to 0.15, where respiration is recruited. **C**. *ε*_*ij*_(P_tot_) of GPM1 (orange), PGK1 (blue) and TDH1 (green) with COX1: line, the COX1 pair; band, the range over all 31 respiratory subunits; open triangles at the right are the FBA values; dotted lines *ε*_*ij*_ = ±0.01.

The groups move in opposite directions. A respiratory-chain or ATP-synthase deletion is neutral while the wild type ferments, because the route it removes then carries no flux in either strain. Once respiration is recruited, the wild type uses that route and the mutant cannot, so the mutant falls to its FBA fitness. Lower-glycolytic deletions do the reverse: harmful while fermentation carries all ATP production, and buffered as respiration takes over. One quantity explains both: the marginal value of protein, *µ*′. In a respiratory deletion it falls to zero at a threshold set by the reaction removed, not by the gene, so the 31 subunits behave as three mutants with three thresholds. The complexes reach their thresholds before the wild type reaches its own, which follows from fermentation being protein-cheap and the respiratory chain not. What the model adds is the distance. Complexes III and IV stop being pool-limited at about a quarter of the wild type’s P_tot_ and ATP synthase at about half, so over most of the physiological window the wild type is still buying protein while these mutants have no use for it. The absolute thresholds shift by more than a factor of three when the turnover numbers are perturbed, but as fractions of the wild type’s they are unchanged, with medians near a quarter and a half and below the wild type’s in every replicate.

The two kinds of respiratory deletion end in different active sets where the pool goes slack. A complex III or IV deletion cannot respire: it ferments and leaves the oxygen bound slack, so glucose alone limits it. An ATP synthase deletion keeps the chain at capacity and saturates the oxygen bound, so glucose and oxygen limit it together. This corresponds to *ϕ* and *δ* phases of Fig 2A, reached by deletion rather than by moving in the phase plane, which is how the P_tot_ sweep produces them while the wild type never enters *ϕ*. Their fitness then falls as P_tot_ rises, only because the wild type keeps growing while they cannot. The lower-glycolytic deletions, PDC1 and the oxidative pentose phosphate entry behave oppositely: their bypass is protein-expensive, so *µ*′ lies above the wild type’s and *W*_*i*_ rises toward one as protein becomes available. A deletion strain thus has its own saturation P_tot_, and by Eq (14) 67 of the 980 viable deletions saturate later than the wild type (Fig D in S1 File), an ordering by how protein-expensive their compensation is.

### The interaction landscape reorganizes with the phases, and one sign inversion is large

We computed all 479,710 pairwise deletions among the 980 viable genes at each of the 421 grid values. Only 2,981 pairs reach |*ε*_*ij*_| ≥ 0.01 at any P_tot_ in the sweep. Approximately 40% of those interact in FBA as well, so their existence owes nothing to the enzyme constraint, and we focus on the remaining 1,741. Fig 3B counts how many of them interact at each P_tot_: few where the pool is the wild type’s only active constraint, with negatives outnumbering positives. Co-limitation reverses the balance: positives climb through the recruitment of respiration until, at the oxygen-saturation boundary, 94% of the pairs the pool creates are alleviating. Most of these come from the 31 respiratory deletions that abolish the marginal value of protein: they take part in more than half of the 1,741 pairs that interact only under the enzyme constraint, four in five of them positive. Above P_tot_ = 0.658 the wild type is FBA and the structure collapses. The pairs that interact in FBA reappear with their FBA sign as the mutants saturate in turn, and the positives that remain at P_tot_ = 0.88 all involve a deletion strain on which the pool constraint is still active, PMT5 and GPM1 among them.

Individual interactions vary smoothly: the largest change in any pair between adjacent grid values is 0.017 (Methods), and no pair interacts with opposite signs at two adjacent values, so a sign change passes through the band below the threshold, |*ε*_*ij*_| *<* 0.01, rather than jumping. The variation is concentrated, however: two in five of all changes in interaction status fall in the 6% of the range over which respiration is recruited, the shaded band of Fig 3B.

Sign inversion is rarer than the reorganization suggests. Of the 2,981 interacting pairs, 596 interact with both signs, and all but 31 are threshold effects: pairs converging on FBA above the pool-slack boundary, respiratory subunits buffering one another once respiration is used, and pairs that never leave the neighborhood of the threshold (Fig E in S1 File). Inside the physiological window, only six change sign at all, and every one is within 0.02 of the threshold. The 31 are one inversion. GPM1 inverts against every subunit of complexes III and IV and of ATP synthase, from −0.22 while the wild type ferments to +0.13 once the wild type respires, and is absent from FBA at both ends (Fig 3C); this is the pair of Fig 1B. Which subunit is lost sets only where the sign crosses, earliest for complex III and latest for ATP synthase.

The two ends have distinct mechanisms. While the wild type ferments, the GPM1 deletion is costly: its bypass through the other phosphoglycerate mutase copies is protein-expensive. The mutant makes up the ATP by respiring while the wild type does not. Removing a respiratory subunit is therefore free in the wild type and a second cost in the mutant. Once the wild type respires, the respiratory deletion takes the cell out of the pool-limited regime, the protein the bypass needs is free, and the double grows as the respiratory single does. The other lower-glycolytic lesions show why GPM1 is the only one that inverts. PGK1, PFK1, PFK2, FBA1 and TPI1 have no cheap bypass, so respiration is conditionally essential for them in FBA as well, and they aggravate with every respiratory subunit at every P_tot_. TDH1 has isozymes that make its deletion cheap, and it buffers throughout and decays to zero. Inversion needs a lesion whose protein-priced cost is relieved by added protein, together with a second lesion whose cost emerges as protein becomes available; in every inverting pair at least one of the two genes is P_tot_-dependent on its own. Closing the three secretion routes is not what produces the inversion: the same 31 pairs invert in ecYeastGEM, and our analysis finds only 2% fewer interacting pairs and 6% fewer P_tot_-only pairs. Under the turnover-number perturbation, the two halves of the inversion separate. The aggravating end is retained in every evaluation where the perturbation preserves the cost of the GPM1 deletion, and it is absent where the perturbation makes the bypass cheap, as the mechanism requires. The crossing to positive epistasis survives in nine of fifteen evaluations at *σ*_*k*_ = 0.3 and three of fifteen at 0.6.

Most of the reorganization is inherited from the single mutants. The 1,741 P_tot_-only pairs involve 126 genes, and 91% of them include at least one of the P_tot_-dependent genes of Fig 3A, as the multiplicative null requires whenever *W*_*i*_ or *W*_*j*_ changes. Co-limitation, not the enzyme constraint as such, produces it: with the pool as the only binding constraint the pair landscape is as P_tot_-invariant as the single mutants (Table 1), and FBA alone gives twice as many interacting pairs as the pool alone.

To assess physiological relevance, we use the window P_tot_ = 0.4 to 0.66, the part of the physiological range in which the pool still binds (Methods). Its upper end is set by our uptake bounds rather than by the cell, and by Eq (11) a higher glucose bound moves it further out. Between its ends, half the landscape is stable and half is not: three in ten of the pairs interacting at either end interact at one end only, and of those interacting at both, three in ten shift by more than 0.05 (Table B in S1 File). The turnover does not depend on where the window sits but scales with its width, falling to one shifted pair between 0.60 and 0.66. The single mutants change less over the same window, none by more than 0.3, and their large transitions lie below P_tot_ = 0.4. Within the physiological range, P_tot_ therefore acts more strongly on interactions than on the single-mutant phenotypes.

### P_tot_-only interactions connect oxidative phosphorylation to the pathways that consume its cofactors, and expose isozymes that FBA cannot see

We grouped the P_tot_-only pairs by the subsystem assignments of their two genes (Methods). Oxidative phosphorylation dominates the network (Fig F in S1 File): the pairs that join it to glycolysis, the citrate cycle, the mitochondrial transporters and the serine, proline and pentose phosphate branches are all enriched over random gene pairs after Bonferroni correction. Enrichment alone says little, since the energy pathways interact with each other in FBA as well, and oxidative phosphorylation with itself far more so. What the pool adds is the connection from oxidative phosphorylation to the biosynthetic branches that draw on its products, serine, proline and the oxidative pentose phosphate pathway, which barely interact in FBA at all.

The two amino-acid branches are almost uniformly positive, and the pentose phosphate pairs split. The three are linked by cofactor traffic: in the model, proline and serine synthesis consume NAD(P)H and the oxidative pentose phosphate pathway supplies NADPH, and the pool constraint charges the enzyme cost of cofactor regeneration to relative fitness. Where the sign is positive, the alleviation is nearly complete: across the positive interactions between oxidative phosphorylation and these branches, a biosynthetic deletion that costs a median 6.8% of growth on its own costs 0.2% once respiration is already deleted. The hubs of the P_tot_-only network are the P_tot_-dependent single mutants, PRO2 and GPM1 foremost. Five genes break the pattern, ACO1 and AQY2 among them, with partners in the dozens while their own fitness varies by less than 0.05 across the sweep.

Holding both single-mutant fitnesses nearly constant would leave no interactions if the reorganization arose entirely from changes in those fitnesses. We applied the strict test of requiring both single-mutant fitnesses to vary by less than 0.005 over the whole sweep, so that no interaction we count can be inherited through the multiplicative null. Thirteen pairs satisfy this condition (Fig 4). All are negative, and nine are model isozymes, alternative gene products of one reaction. Each single deletion is buffered by its isozyme, giving *W*_*i*_ = *W*_*j*_ = 1 at every grid value, and the double removes the reaction. Neutral singles with a harmful double is the signature of synthetic sickness between redundant genes. Redundancy alone does not produce it: FBA has the same alternative routes and none of these pairs interacts there, because a route that costs no enzyme is a backup the cell keeps for free. The isozyme pairs that do interact in FBA are those whose shared reaction is needed, or whose single is itself deleterious.

**Fig 4.**
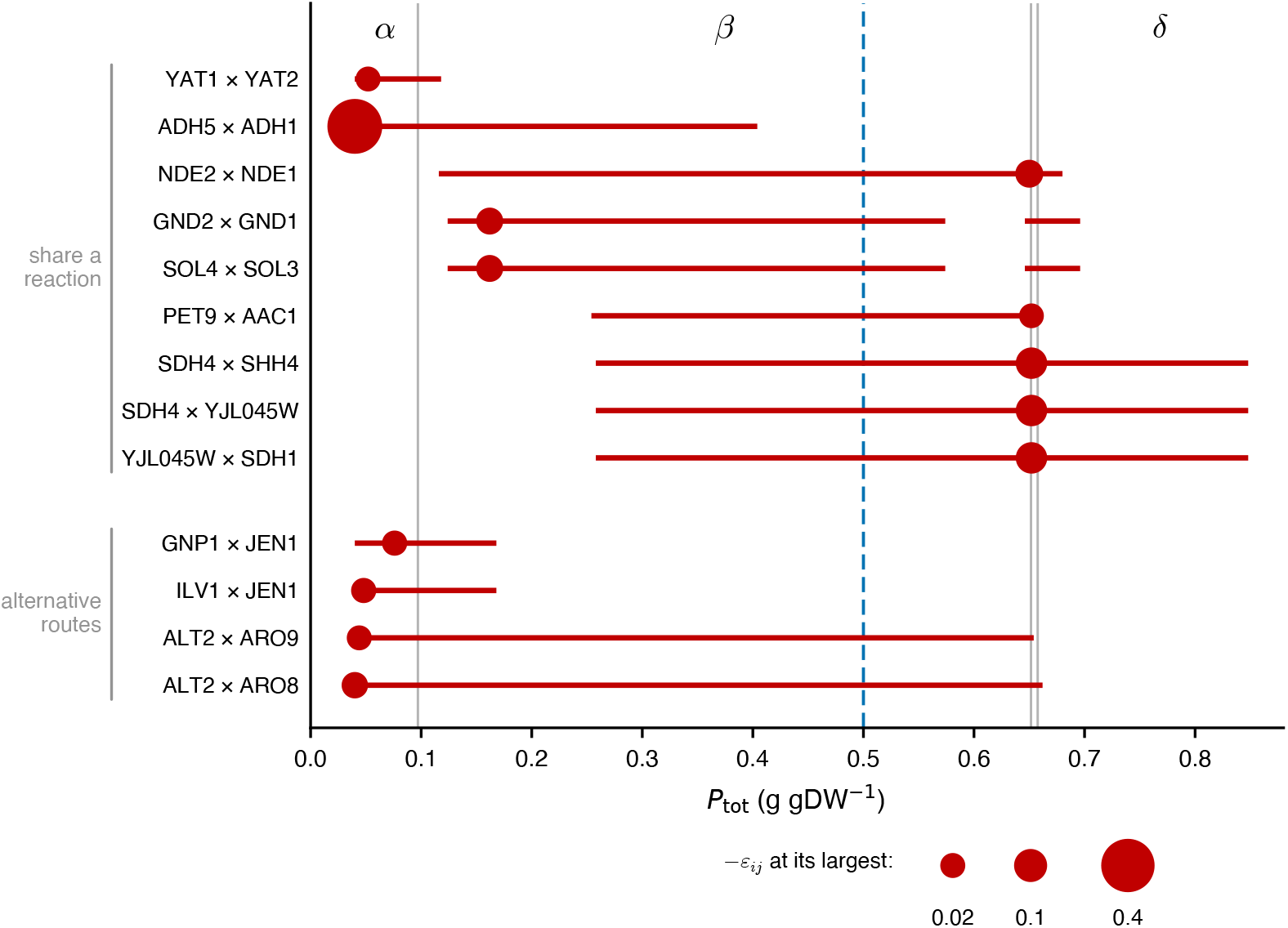
Response of strict pairs (all negative) to varying P_tot_. Red lines, the P_tot_ range over which |*ε*_*ij*_| ≥ 0.01 for a pair; circle, location of the largest interaction, size increasing with− *ε*_*ij*_ (key below). Rows grouped by whether the two genes share a reaction or are alternative routes to one metabolite. Gray verticals and *α, β, δ*, the phases of Fig C in S1 File; dashed blue, the calibration value P_tot_ = 0.5 as in Fig 2A.

Under the pool constraint the alternative route costs more enzyme, and the double mutant pays for it. The pair interacts over the range of P_tot_ in which the shared reaction carries flux (Fig 4). ADH1 with ADH5 is largest at the lowest P_tot_ sampled, *ε*_*ij*_ = −0.43, and gone by 0.40: it lives where fermentation carries the carbon. The pairs of the respiratory machinery appear only once respiration is being recruited and peak just before the pool goes slack. Two of them, GND2 with GND1 and SOL4 with SOL3, fall just under the cut-off between P_tot_ = 0.58 and 0.64, where |*ε*_*ij*_| reaches 0.009, so their rows break in two. We term such an interaction *protein-priced*; it is a network-level effect. The model routes around the two deletions, and the cost arises because the rerouted flux draws enzyme away from the rest of the network. The interaction is therefore a property of the whole flux distribution rather than of the two genes. No single P_tot_ gives all thirteen: at the calibrated 0.5 (dashed blue in Fig 4, as in Fig 2A) nine interact and ADH1 with ADH5 does not.

The remaining four strict pairs share no reaction, but each is a pair of alternative routes to one metabolite, ILV1 and JEN1 for instance, deamination of threonine against import of 2-oxobutanoate, and they sort the same way. The set barely depends on the closed secretion routes, twelve of the thirteen being strict in ecYeastGEM as well, and no strict pair interacts with the opposite sign in any of our turnover-number samples: the interaction shrinks toward the threshold rather than reversing. Network structure shows that these pairs are redundant, and it shows the same in FBA, in which none of them interact. Where two routes exist, FBA is indifferent between them. However with the protein-pool constraint in place, the cheaper route is used, and the more expensive one is a cost that appears only when the first is removed. The interaction therefore needs both ingredients: the topology provides the second route, and the pool gives it a price. This is the pattern documented for duplicate loss in the interaction maps [44].

### Two pathways, one pool: the mechanism at the flux level

To determine the flux-level basis of the two signs, we solved the wild type, both single deletions and the double for a negative and a positive pair on ecYeastGEM-A and followed where the protein goes (Fig 5, Table C in S1 File). The two cases are the two outcomes of competition for one pool: an aggravating interaction, in which the double is worse than the product of its singles, and an alleviating one, in which it is better. In both, the pool is fully used in all four strains, so what changes is allocation, and the four bars of Fig 5B and D have the same length for that reason: what one route gains, another loses. A pair aggravates when the rerouted fluxes of the two deletions draw on the pool at once. It alleviates when one deletion removes the route that the other’s bypass competes with.

**Fig 5.**
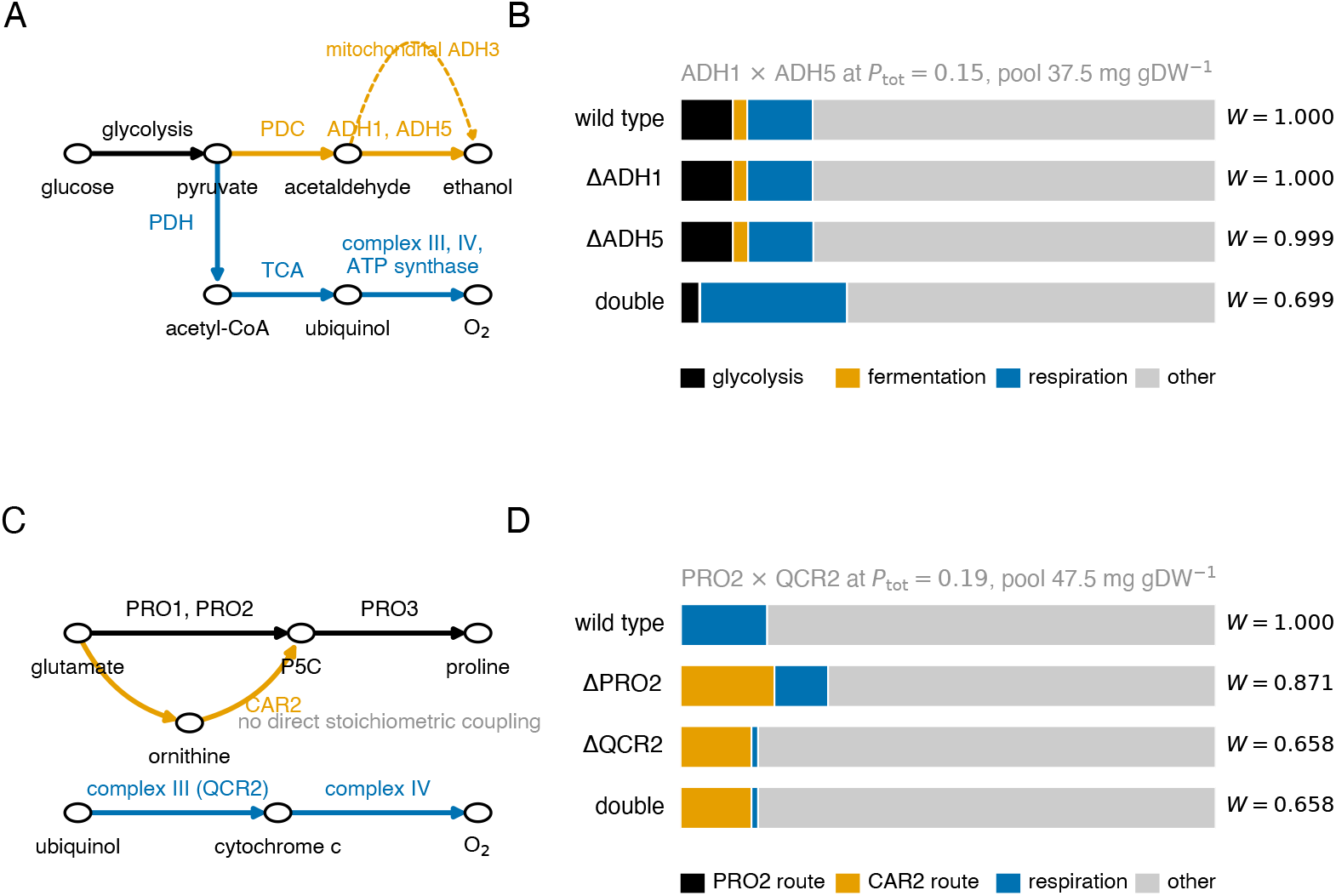
Where the protein goes in a negative and a positive protein-priced interaction. (ecYeastGEM-A). **A**. Routes in ADH1 with ADH5: glycolysis (black), fermentation (orange; dotted, the mitochondrial ADH3) and respiration (blue). **B**. The pool of each genotype at P_tot_ = 0.15, 37.5 mg gDW^−1^ in every bar, divided into the enzyme of each route of **A**; gray, everything else; *W*, relative fitness. **C**. Routes in PRO2 with QCR2: proline through PRO1, PRO2 and PRO3 (black) or through ornithine and CAR2 (orange), and the respiratory chain (blue); the two have no direct stoichiometric coupling. **D**. As **B** at P_tot_ = 0.19, pool 47.5 mg gDW^−1^; the direct proline route, 0.03 mg gDW^−1^, is too small to see.

Consider ADH1 with ADH5 at P_tot_ = 0.15 as the aggravating case. Neither deletion matters on its own, because each isozyme masks the loss of the other, and FBA predicts no interaction for that reason. Remove both and the cytosolic route from acetaldehyde to ethanol closes (Fig 5A), leaving only the mitochondrial alcohol dehydrogenase ADH3, which carries little and cannot carry more. What follows is not a local detour but a change of strategy. Ethanol excretion nearly vanishes, oxygen uptake more than doubles, and the protein committed to the respiratory chain and ATP synthase does the same (Fig 5B). Respiration is so much more expensive per unit of glucose that the cell can no longer use the glucose it is offered: uptake falls well below the bound and the glycolytic enzyme shrinks with it. The cost of losing cytosolic alcohol dehydrogenase is thus distributed across glycolysis and the respiratory chain rather than confined to the deleted reaction, and *ε*_*ij*_ = −0.30. Eq (18) puts the protein the double would need to reach the multiplicative expectation at 41% of P_tot_ itself, a lower bound since *µ*′ falls as P_tot_ rises; the succinate dehydrogenase pairs cost 8% by the same measure.

To assess whether these cases are typical, we solved every interacting pair as a double deletion and recorded the reduced cost of the pool bound. Two numbers came back, and both were larger than we expected. First, the price of Eq (18) is defined at all only where the double is itself pool-limited, and that is a shrinking minority as protein becomes abundant: of the pairs interacting at P_tot_ = 0.15, 0.30 and 0.50, the double retains a positive *µ*′ in just 79%, 53% and 37%. For the rest, PRO2 with QCR2 among them, the second deletion has taken the strain out of the pool-limited regime altogether, and no amount of protein closes the gap. Second, where the price is defined it usually exceeds the protein there is to spend. Among the negative pairs it is larger than P_tot_ itself in 84%, 56% and 58% of cases at those three values, and smaller than a tenth of P_tot_ in at most 14%. Both are first-order extrapolations, and both point the same way. Whatever the pool creates, it does not create marginal effects that a little more enzyme would sweep away.

The alleviating case is less intuitive. PRO2 contributes to proline synthesis, whereas QCR2 is a subunit of complex III; the two have no direct stoichiometric coupling, occupy different compartments and belong to distinct subsystems, yet they interact at P_tot_ = 0.19. Their protein allocation reveals the mechanism (Fig 5C, D). In the wild type PRO2 makes proline at negligible cost. Delete it and proline synthesis is forced through the ornithine route, whose aminotransferase CAR2 demands a sixth of the pool at this availability. That protein comes from respiration: complex III shrinks, ethanol rises, and growth falls. Delete QCR2 instead and the cost is larger, but in that background the optimum already routes proline through CAR2. PRO2 is then no longer used, the second deletion is free, and the double grows exactly as the complex III single does, *ε*_*ij*_ = +0.09. The duals agree: in the PRO2 single the marginal value of protein lies above the wild type’s, the signature of a protein-expensive bypass, while in the complex III single and in the double it is zero, so Eq (18) assigns this interaction no finite price and the alleviation is exact. A deletion is only as costly as the protein its bypass consumes, and that cost is payable only while something else competes for the pool; remove the competitor and the deletion becomes free. Neither interaction is present in FBA, where both bypasses are available at no enzyme cost. The pool therefore places edges in the interaction network where stoichiometry places none, and it does so in both signs.

## Discussion

Four ingredients of this argument are already established. Enzyme constraints have been known to produce overflow metabolism where FBA does not since they were introduced [4, 8]. That the switch requires two growth-limiting constraints is the extremum principle of de Groot and co-workers, and the conditions for the shift have since been derived for a fixed protein content [23–25]. Linearity in the protein constraint is a property of the linear program and has been remarked on for these models more than once: the authors of MOMENT noted that varying the total enzyme mass scales the predicted growth rates and leaves their comparison with measurement unchanged [5], as did Nilsson and Nielsen for yeast [10]. Alzoubi and co-workers noted in passing that their enzyme-mass limit cancels in the ratio of knockout to wild-type biomass flux, so its value does not affect their epistasis predictions [45]. The consequence for genetic interactions had not been drawn. Whenever P_tot_ is the only binding constraint it is a pure scale factor for every mutant as well as for the wild type, up to a maintenance dilution worth at most 0.052 in relative fitness in our screen (Table 1); no interaction changed sign there. That configuration is where enzyme-constrained models are calibrated, and where enzyme constraints have been claimed not to improve the prediction of measured yeast epistasis [45]: in those models the enzyme-mass limit replaces the constraint on nutrient uptake, and with the pool acting alone no relative phenotype can change. The combination that can, a protein constraint binding alongside a nutrient constraint, was not considered.

The effect of P_tot_ on the interaction landscape has two components. Most of what changes is inherited. P_tot_ decides how much respiration the wild type recruits, so it sets the deletion phenotypes of the respiratory machinery and of lower glycolysis. Under the multiplicative null, every interaction involving one of those genes follows them. This component follows from single-mutant phenotypes: the respiratory deletions leave the pool-limited regime, the lower-glycolytic ones pay for a protein-expensive bypass, and where the two kinds of lesion meet we find the one large sign inversion, GPM1 with the respiratory chain. A smaller, structured pairwise component lies beyond this inherited variation: the interactions that exist only under the pool constraint connect oxidative phosphorylation to the biosynthetic branches that consume its cofactors, which FBA misses because it assigns no enzyme cost to regenerating them, and what survives the strictest control, holding both single mutants still, is negative and mostly isozyme pairs, dispensable in FBA and expensive under the pool constraint.

The pool is a constraint of a different reach from stoichiometry. Mass balance ties a reaction to those it shares metabolites with, and through them to their coupled sets [26]; the pool ties every enzyme-catalyzed reaction to every other, at a weight set by its cost per unit flux. The pool is fully used where it binds, so rerouting is zero-sum competition over enzyme, the premise of every proteome-allocation model [46, 47]. The epistasis this produces has two ingredients. The topology of the network decides whether a deletion has an alternative route and whether two routes meet, and the pool decides what each route costs. Two genes therefore need no direct stoichiometric coupling to interact, only competing demands on the pool, and the sign follows from the competition: a pair aggravates when both rerouted fluxes draw on the pool at once, and it alleviates when one deletion removes the route its partner’s bypass competes with, or takes the strain out of the pool-limited regime altogether. This competition occurs wherever the pool binds; only under co-limitation does its outcome depend on the protein available. The same mechanism limits what such a prediction can mean: any enzyme-constrained model with isozyme expansion will predict synthetic sickness between paralogs wherever its turnover numbers make one copy cheaper than the other, and that prediction is only as accurate as those two numbers. That expression costs contribute genetic interactions of both signs has been shown experimentally for a two-gene pathway, where fitness interactions between expression variants were explained quantitatively by flux gains, intermediate toxicity and expression costs [48]. Eq (18) is the same currency read out of a genome-scale model.

Condition-dependent rewiring has experimental precedent. Interaction networks measured under two conditions differ substantially [49], and the epistasis among whole-genome duplicates is extensive and condition-dependent [50]. Those measurements do not, however, identify the governing parameter: which variable shifts the interactions, in which direction and through what mechanism. The framework here addresses one such parameter, protein availability against nutrient supply. A direct comparison with the existing compendia is not discriminating, however (Table D in S1 File). Genome-wide screens are typically performed in batch culture on glucose, the respirofermentative regime in which respiratory genes are repressed, so the model’s interaction pattern there and the regulatory expectation coincide. Eleven of the thirteen strict pairs were measured [51]. Two are scored as interactions at the screen’s own threshold, |*ε*| *>* 0.08 at *p <* 0.05, and both are negative as in the model. The other nine fall below that threshold, and none is scored positive. GPM1 is absent from the screen in every allele, so the sign inversion cannot be tested against it at all. This is consistency, not a test. The optimality premise is best supported in nutrient-limited steady state, the regime of the chemostat physiology above, and the predictions below are therefore contrasts between controlled-supply conditions.

Representing proteome capacity as a single pool has a consequence that compartmentalized models do not share: *D*_crit_ scales linearly with protein availability (Eq (11)). pcYeast and a resource-balance model of yeast attribute the Crabtree threshold instead to the capacity of the inner mitochondrial membrane [52, 53], and one experiment favors them: a heterologous alternative oxidase raised *D*_crit_ of a CEN.PK strain by 10% [41], whereas adding that oxidase to ecYeastGEM-A at any enzyme cost leaves *D*_crit_ unchanged, because at the transition the optimum is already the maximum-yield respiratory mode. A compartment-specific pool is the natural repair, and we have not fitted one, since its capacity would be a free parameter calibrated on the transition it is meant to explain. The homogeneity argument extends to it unchanged: a second pool acts on relative phenotypes only through its ratio to the first.

The sign structure also separates the species. A Crabtree-negative yeast does not cross to the fermentative strategy under these conditions, so the glycolysis–respiration inversion should be absent from its interaction landscape. An engineered Crabtree-negative *S. cerevisiae* [43] characterized with the same enzyme-constrained model, is a good candidate strain in which to look.

Three considerations delimit the interpretation. First, the adjustment we found necessary reveals a limitation of the model. Away from its calibration point, ecYeastGEM prefers non-physiological secretions by a fraction of a percent in growth rate, a near-tie resolved by the solver rather than by biology, so any study that leaves the calibration configuration should check what the model excretes. The closures change the landscape little; what they decide is which regime it describes. The group that built pcYeast reports the same kind of near-tie: the relative proteome efficiency of respiration and fermentation shifts within a 5% range of the turnover numbers used, and their model switches strategy under similarly small changes [54]. In this region of parameter space, a model’s choice of strategy is set by parameters that are not known to that precision.

Second, the turnover numbers are heterogeneous in provenance [9], and the results are robust to them in structure but not in membership. The P_tot_-dependent set, the sign of every strict pair, and the aggravating phase of the inversion survives a lognormal perturbation of every *k*^cat^; the crossing to positive does not. Individual pairs are predictions of this parameter set, and the contrasts between conditions are the robust part.

Third, every prediction presented here assumes optimal growth. That is defensible for a wild type shaped by selection and less so for a freshly constructed deletion strain, which is the observation that motivated MOMA and ROOM [55, 56]. Whether the contrasts between conditions survive under non-optimal objectives remains to be tested.

Four contrasts follow that experiments can test. First, by Eq (11) the region *β*, where the pool binds alongside the glucose bound, widens in proportion to the glucose uptake. At high uptake *β* covers the whole physiological range of protein availabilities. At low uptake it covers none of that range: in a glucose-limited chemostat below *D*_crit_ the pool is slack, and the wild type solves the FBA problem. Interactions among the genes identified here should therefore be condition-dependent above *D*_crit_ and absent below it, a contrast that condition-resolved screens [22] can test. By Eq (10), the dependence of interactions on glucose supply at constant proteome is the direct experimental counterpart of the P_tot_ dependence computed here. Second, sign inversion between growth regimes follows for one specific kind of pair. It is a glycolytic lesion whose cost is relieved by enzyme, such as a hypomorph or a deletion backed by a weaker isozyme, together with a respiratory deletion. Such a pair should aggravate in fermentative growth, where the lesion makes respiration conditionally essential, and buffer in respiratory growth, where the respiratory deletion frees the protein the lesion needs. GPM1 is the model’s example, and lesions without a backup, PGK1 or PFK1, should aggravate in both regimes. A perturbation that lowers the protein available to metabolism, a proteome burden or a stress response, pushes the cell toward the fermentative side of the boundary and should carry these signs with it. Third, the respiratory-phase isozyme pairs whose shared reaction is dispensable in FBA should be synthetic-sick where the pool constraint limits respiration, in respirofermentative growth, and neutral under glucose limitation. These are the corresponding pairs in Fig 4, the succinate dehydrogenase compositions foremost, and they span the physiological window. The discriminating observation is their predicted disappearance under glucose limitation. Fourth, our analysis suggests that interactions between the respiratory chain and serine, proline and oxidative pentose phosphate biosynthesis appear when protein availability is tightened, by stress or by heterologous expression, and disappear when it is relaxed. They should be positive for the amino-acid branches and mostly negative for the pentose phosphate pairs.

## Author contributions

Conceptualization, methodology, software, formal analysis, investigation, visualization, writing and supervision: EA. Investigation in an earlier phase of the study: TK. Both authors read and approved the final manuscript.

## Supporting information

**S1 File. Supporting figures and tables**. Figs A to F and Tables A to D.

## Notes

https://doi.org/10.5281/zenodo.22705816

https://github.com/AlmaasLab/ecFBA-epistasis

### Competing Interest Statement

The authors have declared no competing interest.

## References

1. Orth JD, Thiele I, Palsson BØ. What is flux balance analysis? Nat Biotechnol. 2010;28(3):245–8. doi:10.1038/nbt.1614.

2. Bordbar A, Monk JM, King ZA, Palsson BO. Constraint-based models predict metabolic and associated cellular functions. Nat Rev Genet. 2014;15(2):107–20. doi:10.1038/nrg3643.

3. Beg QK, Vazquez A, Ernst J, de Menezes MA, Bar-Joseph Z, Barabási AL, et al. Intracellular crowding defines the mode and sequence of substrate uptake by Escherichia coli and constrains its metabolic activity. Proc Natl Acad Sci USA. 2007;104(31):12663–8. doi:10.1073/pnas.0609845104.

4. Vazquez A, Beg QK, deMenezes MA, Ernst J, Bar-Joseph Z, Barabási AL, et al. Impact of the solvent capacity constraint on E. coli metabolism. BMC Syst Biol. 2008;2:7. doi:10.1186/1752-0509-2-7.

5. Adadi R, Volkmer B, Milo R, Heinemann M, Shlomi T. Prediction of microbial growth rate versus biomass yield by a metabolic network with kinetic parameters. PLoS Comput Biol. 2012;8(7):e1002575. doi:10.1371/journal.pcbi.1002575.

6. Bekiaris PS, Klamt S. Automatic construction of metabolic models with enzyme constraints. BMC Bioinformatics. 2020;21(1):19. doi:10.1186/s12859-019-3329-9.

7. Mori M, Hwa T, Martin OC, De Martino A, Marinari E. Constrained allocation flux balance analysis. PLoS Comput Biol. 2016;12(6):e1004913. doi:10.1371/journal.pcbi.1004913.

8. Sánchez BJ, Zhang C, Nilsson A, Lahtvee PJ, Kerkhoven EJ, Nielsen J. Improving the phenotype predictions of a yeast genome-scale metabolic model by incorporating enzymatic constraints. Mol Syst Biol. 2017;13(8):935. doi:10.15252/msb.20167411.

9. Chen Y, Gustafsson J, Tafur Rangel A, Anton M, Domenzain I, Kittikunapong C, et al. Reconstruction, simulation and analysis of enzyme-constrained metabolic models using GECKO Toolbox 3.0. Nat Protoc. 2024;19(3):629–67. doi:10.1038/s41596-023-00931-7.

10. Nilsson A, Nielsen J. Metabolic trade-offs in yeast are caused by F1F0-ATP synthase. Sci Rep. 2016;6:22264. doi:10.1038/srep22264.

11. Chen Y, Nielsen J. Energy metabolism controls phenotypes by protein efficiency and allocation. Proc Natl Acad Sci U S A. 2019;116(35):17592–7. doi:10.1073/pnas.1906569116.

12. van Hoek P, van Dijken JP, Pronk JT. Effect of specific growth rate on fermentative capacity of baker’s yeast. Appl Environ Microbiol. 1998;64(11):4226–33. doi:10.1128/AEM.64.11.4226-4233.1998.

13. Xia J, Sánchez BJ, Chen Y, Campbell K, Kasvandik S, Nielsen J. Proteome allocations change linearly with the specific growth rate of Saccharomyces cerevisiae under glucose limitation. Nat Commun. 2022;13(1):2819. doi:10.1038/s41467-022-30513-2.

14. Björkeroth J, Campbell K, Malina C, Yu R, Di Bartolomeo F, Nielsen J. Proteome reallocation from amino acid biosynthesis to ribosomes enables yeast to grow faster in rich media. Proc Natl Acad Sci U S A. 2020;117(35):21804–12. doi:10.1073/pnas.1921890117.

15. Yu R, Campbell K, Pereira R, Björkeroth J, Qi Q, Vorontsov E, et al. Nitrogen limitation reveals large reserves in metabolic and translational capacities of yeast. Nat Commun. 2020;11(1):1881. doi:10.1038/s41467-020-15749-0.

16. Gasch AP, Spellman PT, Kao CM, Carmel-Harel O, Eisen MB, Storz G, et al. Genomic expression programs in the response of yeast cells to environmental changes. Mol Biol Cell. 2000;11(12):4241–57. doi:10.1091/mbc.11.12.4241.

17. Metzl-Raz E, Kafri M, Yaakov G, Soifer I, Gurvich Y, Barkai N. Principles of cellular resource allocation revealed by condition-dependent proteome profiling. Elife. 2017;6:e28034. doi:10.7554/eLife.28034.

18. Segrè D, DeLuna A, Church GM, Kishony R. Modular epistasis in yeast metabolism. Nat Genet. 2005;37(1):77–83. doi:10.1038/ng1489.

19. He X, Qian W, Wang Z, Li Y, Zhang J. Prevalent positive epistasis in Escherichia coli and Saccharomyces cerevisiae metabolic networks. Nat Genet. 2010;42(3):272–6. doi:10.1038/ng.524.

20. Kryazhimskiy S. Emergence and propagation of epistasis in metabolic networks. Elife. 2021;10:e60200. doi:10.7554/eLife.60200.

21. Snitkin ES, Segrè D. Epistatic interaction maps relative to multiple metabolic phenotypes. PLoS Genet. 2011;7(2):e1001294. doi:10.1371/journal.pgen.1001294.

22. Costanzo M, Hou J, Messier V, Nelson J, Rahman M, VanderSluis B, et al. Environmental robustness of the global yeast genetic interaction network. Science. 2021;372(6542):eabf8424. doi:10.1126/science.abf8424.

23. de Groot DH, van Boxtel C, Planqué R, Bruggeman FJ, Teusink B. The number of active metabolic pathways is bounded by the number of cellular constraints at maximal metabolic rates. PLoS Comput Biol. 2019;15(3):e1006858. doi:10.1371/journal.pcbi.1006858.

24. de Groot DH, Lischke J, Muolo R, Planqué R, Bruggeman FJ, Teusink B. The common message of constraint-based optimization approaches: overflow metabolism is caused by two growth-limiting constraints. Cell Mol Life Sci. 2020;77(3):441–53. doi:10.1007/s00018-019-03380-2.

25. Droste MJ, Planqué R, Bruggeman FJ. On the conditions for shifts in metabolic strategies. PLoS Comput Biol. 2026;22(6):e1014417. doi:10.1371/journal.pcbi.1014417.

26. Burgard AP, Nikolaev EV, Schilling CH, Maranas CD. Flux coupling analysis of genome-scale metabolic network reconstructions. Genome Res. 2004;14(2):301–12. doi:10.1101/gr.1926504.

27. Müller S, Regensburger G, Steuer R. Enzyme allocation problems in kinetic metabolic networks: optimal solutions are elementary flux modes. J Theor Biol. 2014;347:182–90. doi:10.1016/j.jtbi.2013.11.015.

28. Wortel MT, Peters H, Hulshof J, Teusink B, Bruggeman FJ. Metabolic states with maximal specific rate carry flux through an elementary flux mode. FEBS J. 2014;281(6):1547–55. doi:10.1111/febs.12722.

29. van den Bogaard S, Saa PA, Alter TB. Sensitivities in protein allocation models reveal distribution of metabolic capacity and flux control. Bioinformatics. 2024;40(12):btae691. doi:10.1093/bioinformatics/btae691.

30. Dourado H, Liebermeister W, Ebenhöh O, Lercher MJ. Mathematical properties of optimal fluxes in cellular reaction networks at balanced growth. PLoS Comput Biol. 2023;19(6):e1011156. doi:10.1371/journal.pcbi.1011156.

31. Lu H, Li F, Sánchez BJ, Zhu Z, Li G, Domenzain I, et al. A consensus S. cerevisiae metabolic model Yeast8 and its ecosystem for comprehensively probing cellular metabolism. Nat Commun. 2019;10(1):3586. doi:10.1038/s41467-019-11581-3.

32. Domenzain I, Sánchez B, Anton M, Kerkhoven EJ, Millán-Oropeza A, Henry C, et al. Reconstruction of a catalogue of genome-scale metabolic models with enzymatic constraints using GECKO 2.0. Nat Commun. 2022;13(1):3766. doi:10.1038/s41467-022-31421-1.

33. Stanley GA, Pamment NB. Transport and intracellular accumulation of acetaldehyde in Saccharomyces cerevisiae. Biotechnol Bioeng. 1993;42(1):24–9. doi:10.1002/bit.260420104.

34. Aranda A, del Olmo M. Exposure of Saccharomyces cerevisiae to acetaldehyde induces sulfur amino acid metabolism and polyamine transporter genes, which depend on Met4p and Haa1p transcription factors, respectively. Appl Environ Microbiol. 2004;70(4):1913–22. doi:10.1128/AEM.70.4.1913-1922.2004.

35. González E, Fernández MR, Larroy C, Solá L, Pericàs MA, Parés X, et al. Characterization of a (2R,3R)-2,3-butanediol dehydrogenase as the Saccharomyces cerevisiae YAL060W gene product. J Biol Chem. 2000;275(46):35876–85. doi:10.1074/jbc.M003035200.

36. Romano P, Suzzi G. Origin and production of acetoin during wine yeast fermentation. Appl Environ Microbiol. 1996;62(2):309–15. doi:10.1128/aem.62.2.309-315.1996.

37. Kim SJ, Seo SO, Jin YS, Seo JH. Production of 2,3-butanediol by engineered Saccharomyces cerevisiae. Bioresour Technol. 2013;146:274–81. doi:10.1016/j.biortech.2013.07.081.

38. Lundin M, Baltscheffsky H, Ronne H. Yeast PPA2 gene encodes a mitochondrial inorganic pyrophosphatase that is essential for mitochondrial function. J Biol Chem. 1991;266(19):12168–72. doi:10.1016/S0021-9258(18)98875-7.

39. Reznik E, Mehta P, Segrè D. Flux imbalance analysis and the sensitivity of cellular growth to changes in metabolite pools. PLoS Comput Biol. 2013;9(8):e1003195. doi:10.1371/journal.pcbi.1003195.

40. Postma E, Verduyn C, Scheffers WA, van Dijken JP. Enzymic analysis of the Crabtree effect in glucose-limited chemostat cultures of Saccharomyces cerevisiae. Appl Environ Microbiol. 1989;55(2):468–77. doi:10.1128/aem.55.2.468-477.1989.

41. Vemuri GN, Eiteman MA, McEwen JE, Olsson L, Nielsen J. Increasing NADH oxidation reduces overflow metabolism in Saccharomyces cerevisiae. Proc Natl Acad Sci U S A. 2007;104(7):2402–7. doi:10.1073/pnas.0607469104.

42. Edwards JS, Ramakrishna R, Palsson BO. Characterizing the metabolic phenotype: a phenotype phase plane analysis. Biotechnol Bioeng. 2002;77(1):27–36. doi:10.1002/bit.10047.

43. Tafur Rangel A, Castillo García A, Malina C, Siewers V, Kerkhoven EJ. Model-driven analysis reveals oxidative stress adaptation enabling efficient energy utilization in a Crabtree-negative Saccharomyces cerevisiae. Sci Rep. 2026. doi:10.1038/s41598-026-58495-x.

44. Ihmels J, Collins SR, Schuldiner M, Krogan NJ, Weissman JS. Backup without redundancy: genetic interactions reveal the cost of duplicate gene loss. Mol Syst Biol. 2007;3:86. doi:10.1038/msb4100127.

45. Alzoubi D, Desouki AA, Lercher MJ. Flux balance analysis with or without molecular crowding fails to predict two thirds of experimentally observed epistasis in yeast. Sci Rep. 2019;9(1):11837. doi:10.1038/s41598-019-47935-6.

46. Molenaar D, van Berlo R, de Ridder D, Teusink B. Shifts in growth strategies reflect tradeoffs in cellular economics. Mol Syst Biol. 2009;5:323. doi:10.1038/msb.2009.82.

47. Basan M, Hui S, Okano H, Zhang Z, Shen Y, Williamson JR, et al. Overflow metabolism in Escherichia coli results from efficient proteome allocation. Nature. 2015;528(7580):99–104. doi:10.1038/nature15765.

48. Kemble H, Eisenhauer C, Couce A, Chapron A, Magnan M, Gautier G, et al. Flux, toxicity, and expression costs generate complex genetic interactions in a metabolic pathway. Sci Adv. 2020;6(23):eabb2236. doi:10.1126/sciadv.abb2236.

49. Bandyopadhyay S, Mehta M, Kuo D, Sung MK, Chuang R, Jaehnig EJ, et al. Rewiring of genetic networks in response to DNA damage. Science. 2010;330(6009):1385–9. doi:10.1126/science.1195618.

50. Musso G, Costanzo M, Huangfu M, Smith AM, Paw J, San Luis BJ, et al. The extensive and condition-dependent nature of epistasis among whole-genome duplicates in yeast. Genome Res. 2008;18(7):1092–9. doi:10.1101/gr.076174.108.

51. Costanzo M, VanderSluis B, Koch EN, Baryshnikova A, Pons C, Tan G, et al. A global genetic interaction network maps a wiring diagram of cellular function. Science. 2016;353(6306):aaf1420. doi:10.1126/science.aaf1420.

52. Elsemman IE, Rodriguez Prado A, Grigaitis P, Garcia Albornoz M, Harman V, Holman SW, et al. Whole-cell modeling in yeast predicts compartment-specific proteome constraints that drive metabolic strategies. Nat Commun. 2022;13(1):801. doi:10.1038/s41467-022-28467-6.

53. Dinh HV, Maranas CD. Evaluating proteome allocation of Saccharomyces cerevisiae phenotypes with resource balance analysis. Metab Eng. 2023;77:242–55. doi:10.1016/j.ymben.2023.04.009.

54. Battjes J, Grigaitis P, Hoving M, Visser TD, Stampfl K, Miguel GA, et al. Mitochondrial efficiency determines Crabtree effect across yeasts. bioRxiv. 2024. Preprint, 10.1101/2024.11.01.621473; no journal version as of September 2026.

55. Segré D, Vitkup D, Church GM. Analysis of optimality in natural and perturbed metabolic networks. Proc Natl Acad Sci U S A. 2002;99(23):15112–7. doi:10.1073/pnas.232349399.

56. Shlomi T, Berkman O, Ruppin E. Regulatory on/off minimization of metabolic flux changes after genetic perturbations. Proc Natl Acad Sci U S A. 2005;102(21):7695–700. doi:10.1073/pnas.0406346102.

