## Supplemental File for "Competing constraints on protein availability and nutrient uptake reshape yeast genetic interactions"

### S1 File. Supporting figures and tables

for

Figs A to F and Tables A to D. Equation and figure numbers refer to the main text; ecYeastGEM-A and the constraint configurations are defined in its Methods.

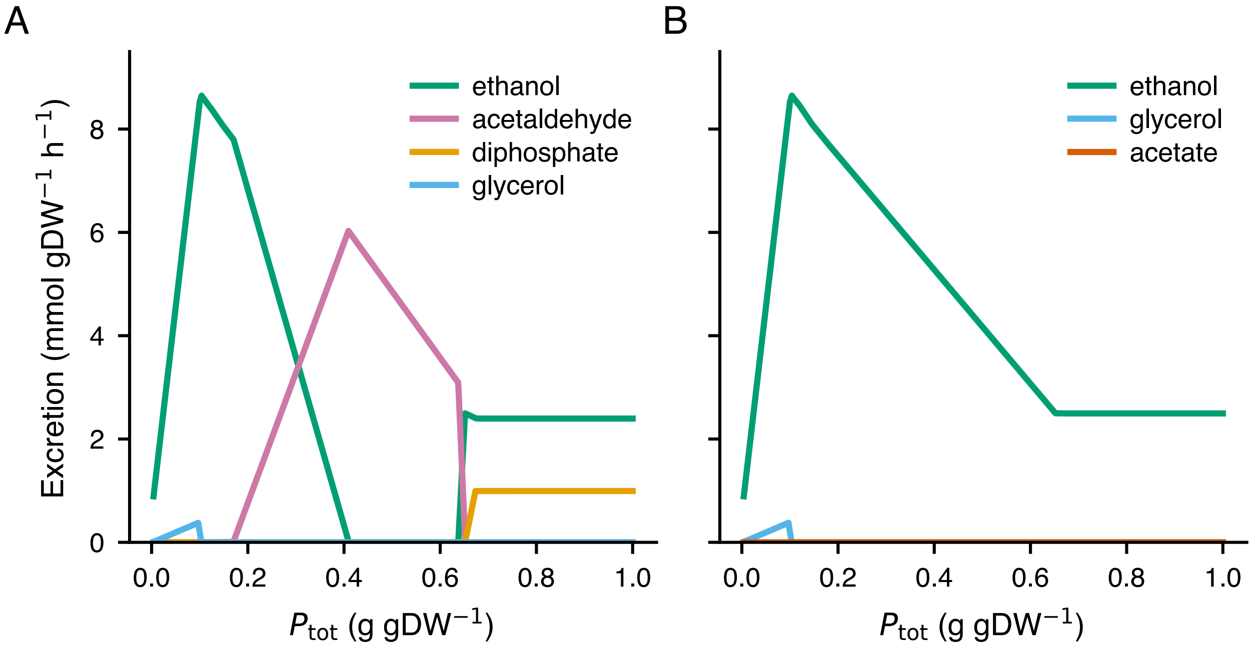


**Fig A. Closing the secretion routes.** Excretion fluxes along the $P_{tot}$ sweep of the main text (glucose bound 5, oxygen bound 8 mmol gDW$^{-1}$ h$^{-1}$) for **A.** ecYeastGEM as published and **B.** ecYeastGEM-A, with the exports of acetaldehyde, (R,R)-2,3-butanediol and diphosphate closed (Methods). As published, the model excretes these three in place of ethanol over most of the range; with them closed, ethanol is the sole overflow product, and growth changes by at most 2%.

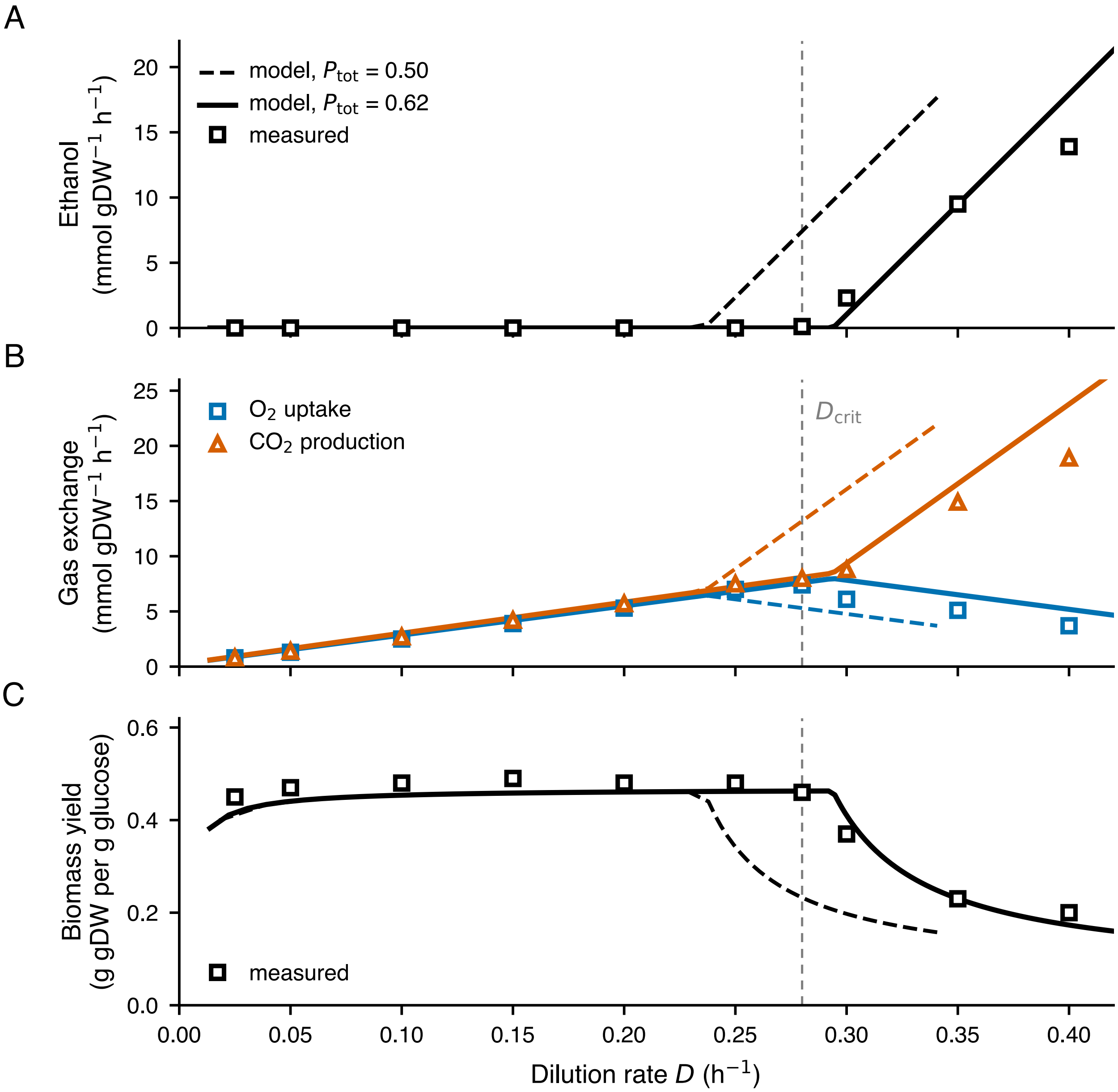


**Fig B. ecYeastGEM-A reproduces glucose-limited chemostat physiology.** Model at the fitted $P_{tot} = 0.62$ g gDW$^{-1}$ (solid lines) and at the published $P_{tot} = 0.50$ (dashed), oxygen unconstrained and the glucose bound scanned, against measurements for *S. cerevisiae* DS28911, Table 1 of [1] (open symbols). **A.** Ethanol production. **B.** Oxygen uptake and carbon dioxide production. **C.** Biomass yield on glucose. Vertical line, the measured $D_{crit} = 0.28$ h$^{-1}$. The fitted value is the least-squares estimate of $P_{tot}$ over all forty measurements (Methods); the published value is drawn for contrast, and nothing else was fitted.

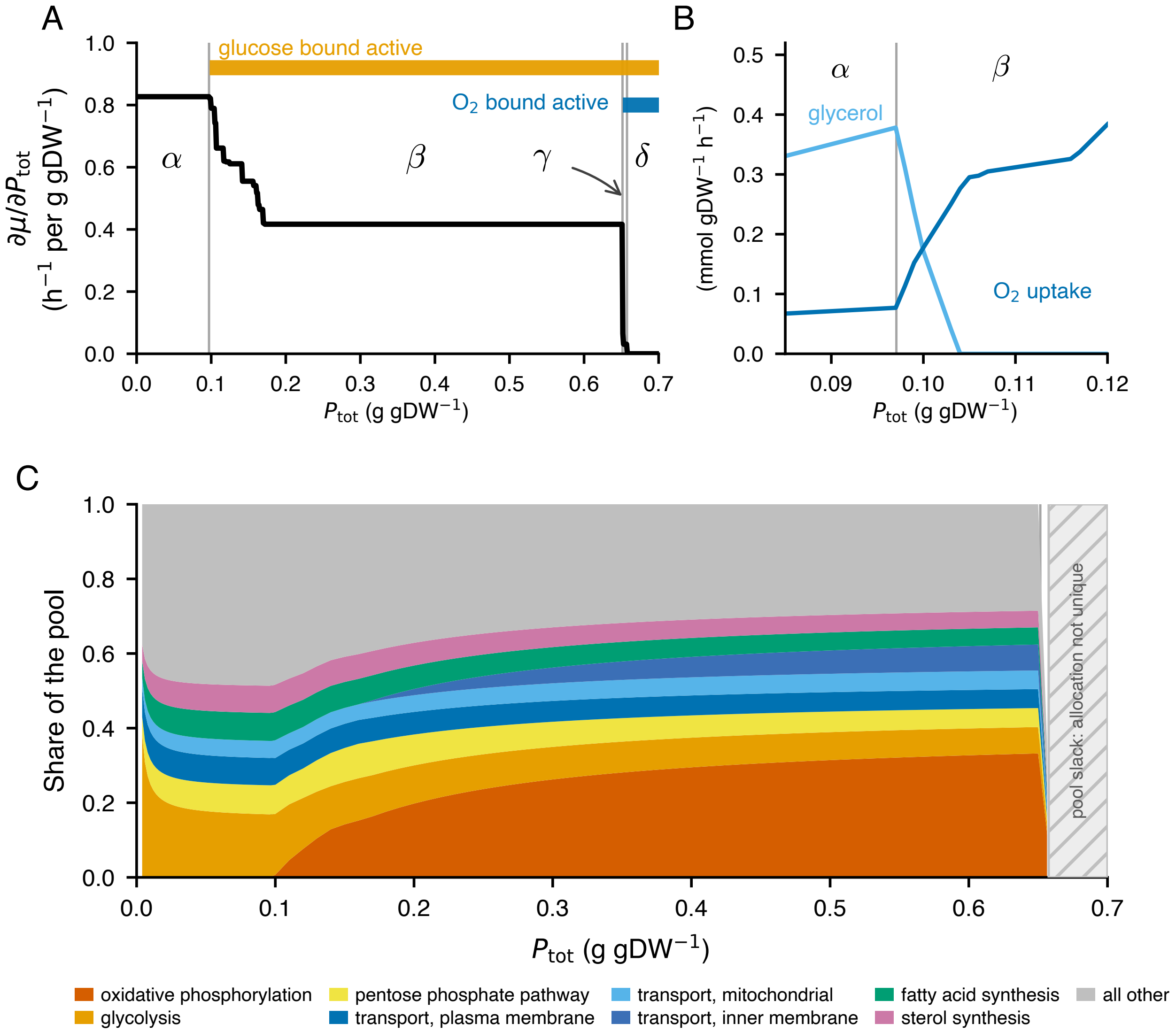


**Fig C. Key phenotypes of the wild type along the $P_{tot}$ sweep** (ecYeastGEM-A, glucose bound 5, oxygen bound 8 mmol $gDW^{-1}$ $h^{-1}$; the red dashed line of Fig 2A of the main text). Gray rules, the active-set changes at $P_{tot} = 0.097$, 0.652 and 0.658; $\alpha$ to $\delta$, the regimes of Fig 2A of the main text along $\nu_{glc} = 5$, the narrow $\gamma$ arrowed. **A.** The marginal value of protein, and which bounds are active. **B.** Glycerol and oxygen exchange across the boundary between $\alpha$ and $\beta$. **C.** Composition of the protein pool by subsystem; hatched, where the pool is slack and the allocation is not unique.

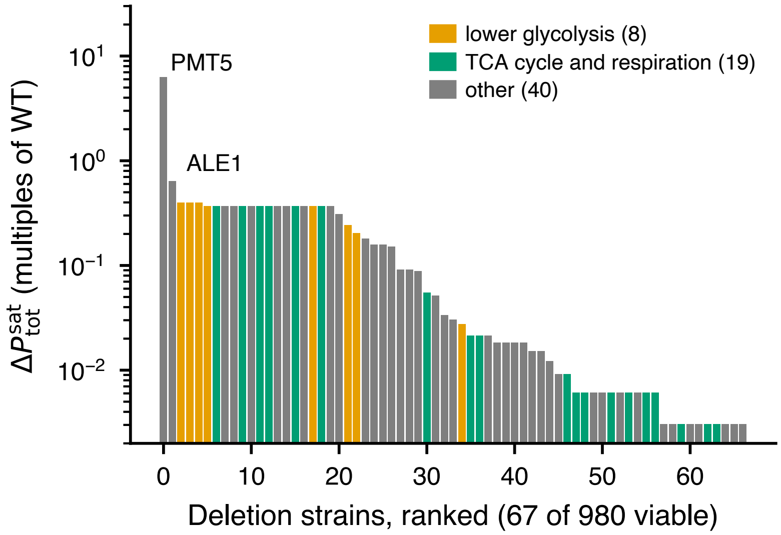


**Fig D. Protein a deletion strain needs before its growth saturates.** $\Delta P_{tot}^{sat}$ of Eq (14) of the main text, in multiples of the wild-type saturation value, for the 67 of 980 viable deletions that saturate after the wild type, ranked on a logarithmic axis (ecYeastGEM-A; the wild type saturates at $P_{tot} = 0.658$, the oxygen-saturation boundary). Orange, lower glycolysis; green, the citrate cycle and the respiratory chain; gray, all others; the two extremes, PMT5 and ALE1, are named. The flat tail is the resolution of $P_{tot}^{sat}$: those strains saturate within 0.002 in $P_{tot}$ of the wild type, the spacing of the sweep there.

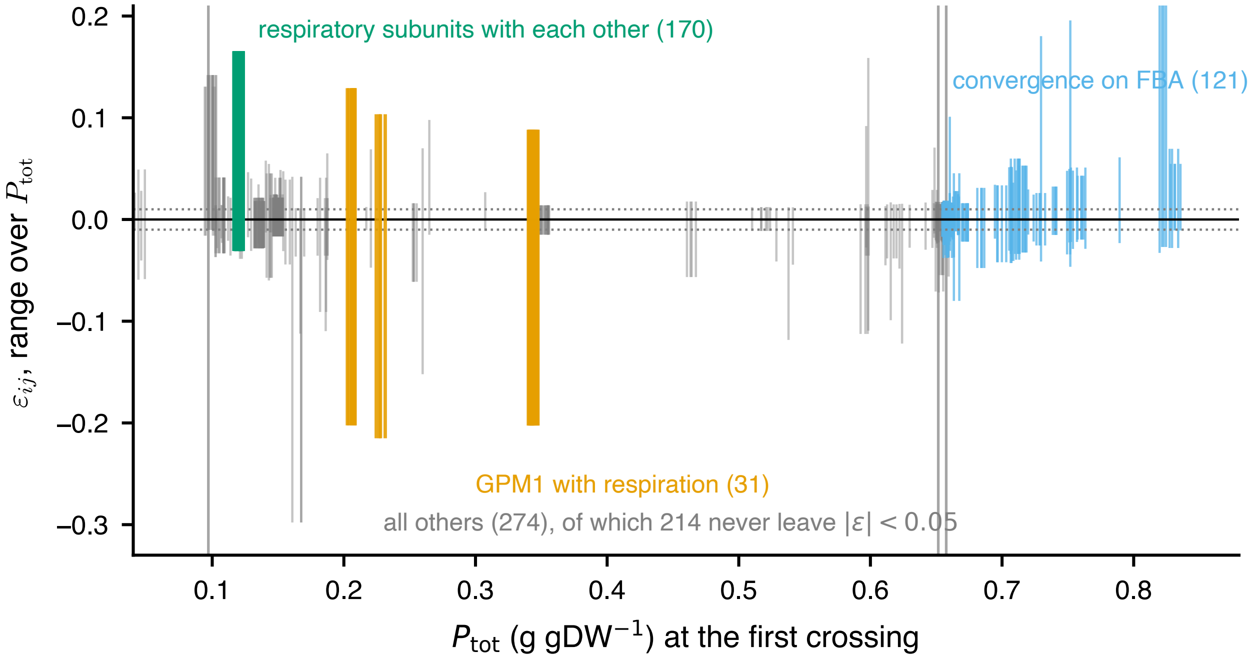


**Fig E. Where the sign-changing pairs change sign, and by how much.** Each of the 596 pairs that interact with both signs across the sweep is drawn as a vertical segment from its most negative to its most positive $\varepsilon_{ij}$ above the threshold, placed at the $P_{tot}$ of its first sign change (Methods; ecYeastGEM-A, interaction threshold $|\varepsilon_{ij}| \geq 0.01$). Orange, GPM1 with a subunit of complexes III and IV or of ATP synthase; green, two such subunits with each other; blue, pairs whose every sign change lies above the pool-slack boundary, where convergence on FBA drives them through the threshold; gray, every other pair. Gray verticals, the active-set changes of Fig C; dotted horizontals, the interaction threshold, $\varepsilon_{ij} = \pm 0.01$.

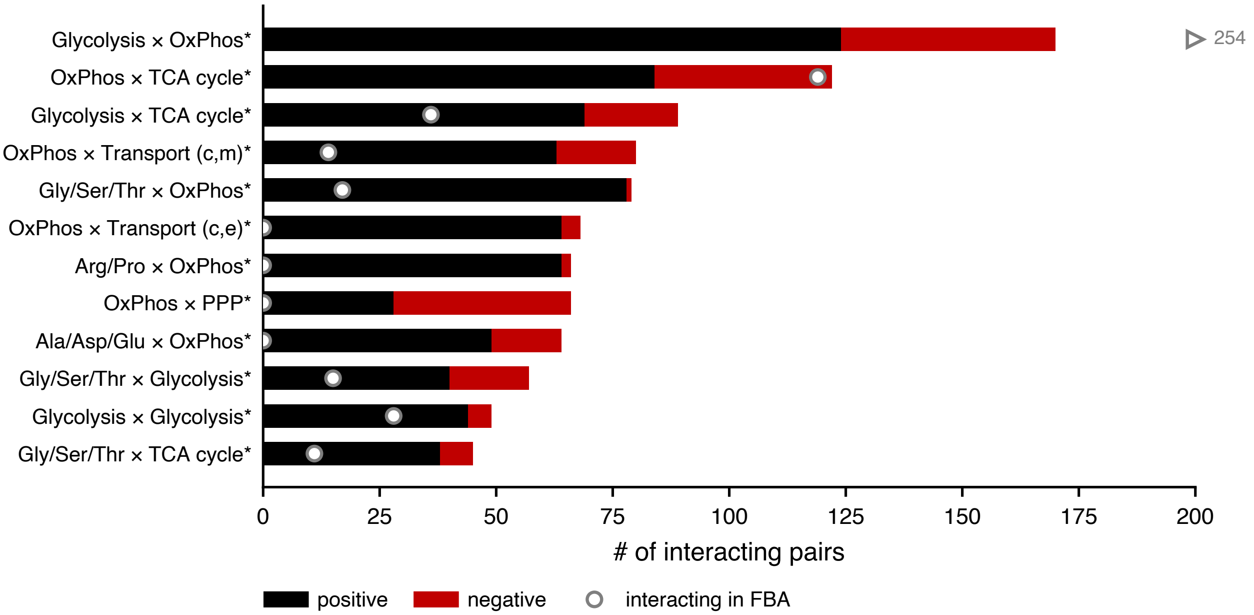


**Fig F. Subsystem structure of the $P_{tot}$-only interactions** (ecYeastGEM-A). The twelve subsystem pairs with the most $P_{tot}$-only interactions (Methods): bar, $P_{tot}$-only pairs, positive black and negative red; open circle, pairs that interact in FBA between the same subsystems, arrowed with the count where off the axis; asterisk, Bonferroni-corrected enrichment $p < 0.05$. Enrichment is relative to random gene pairs among the 980 screened genes (Methods); the energy pathways interact in FBA as well, whereas the connections from oxidative phosphorylation to the serine, proline and pentose phosphate branches are almost absent there.

**Table A. Measured and predicted physiology of aerobic glucose-limited chemostats.** Measurements for *S. cerevisiae* DS28911 from Table 1 of van Hoek and co-workers [1]; model, ecYeastGEM-A at $P_{tot} = 0.62$ g gDW$^{-1}$ with oxygen unconstrained and the glucose bound scanned, interpolated at the measured dilution rates. Specific rates $\nu$ in mmol gDW$^{-1}$ h$^{-1}$, biomass yield $Y_{sx}$ in g biomass per g glucose.

| | $\nu_{\text{ethanol}}$ | | $\nu_{O_2}$ | | $\nu_{CO_2}$ | | $Y_{sx}$ | |
|---|---|---|---|---|---|---|---|---|
| $D$ (h$^{-1}$) | meas. | model | meas. | model | meas. | model | meas. | model |
| 0.025 | 0 | 0 | 0.8 | 0.9 | 0.8 | 0.9 | 0.45 | 0.42 |
| 0.05 | 0 | 0 | 1.3 | 1.5 | 1.4 | 1.6 | 0.47 | 0.44 |
| 0.10 | 0 | 0 | 2.5 | 2.9 | 2.7 | 3.0 | 0.48 | 0.45 |
| 0.15 | 0 | 0 | 3.9 | 4.2 | 4.2 | 4.4 | 0.49 | 0.46 |
| 0.20 | 0 | 0 | 5.3 | 5.5 | 5.7 | 5.8 | 0.48 | 0.46 |
| 0.25 | 0 | 0 | 7.0 | 6.8 | 7.5 | 7.2 | 0.48 | 0.46 |
| 0.28 | 0.11 | 0 | 7.4 | 7.6 | 8.0 | 8.1 | 0.46 | 0.46 |
| 0.30 | 2.3 | 1.0 | 6.1 | 7.8 | 8.8 | 9.4 | 0.37 | 0.41 |
| 0.35 | 9.5 | 9.5 | 5.1 | 6.5 | 14.9 | 16.6 | 0.23 | 0.23 |
| 0.40 | 13.9 | 17.9 | 3.7 | 5.2 | 18.9 | 23.8 | 0.20 | 0.17 |

**Table B. Turnover of the interaction landscape between two $P_{tot}$ values, for four choices of window** (ecYeastGEM-A; interacting, $|\varepsilon| \geq 0.01$; shifted, $|\Delta\varepsilon| \geq 0.05$ among pairs interacting at both). First row, the physiological window of the main text.

| $P_{tot}$ pair | interacting at first | interacting at second | at both | shifted |
|---|---|---|---|---|
| 0.40 vs 0.66 | 1863 | 1973 | 1580 | 456 |
| 0.30 vs 0.60 | 1705 | 2078 | 1543 | 486 |
| 0.50 vs 0.66 | 1971 | 1973 | 1731 | 148 |
| 0.60 vs 0.66 | 2078 | 1973 | 1866 | 1 |

**Table C. Flux-level mechanism of a negative and a positive $P_{tot}$-only interaction** (ecYeastGEM-A). Uptake and excretion in mmol $gDW^{-1}$ $h^{-1}$; protein committed to complex III, summed over its nine subunits, and to CAR2, in mg $gDW^{-1}$. The pool is fully used in every genotype, so what changes between them is allocation.

| *ADH1* × *ADH5 at* $P_{tot} = 0.15$; $\varepsilon_{ij} = -0.30$ | | | | |
|---|---|---|---|---|
| | wild type | ΔADH1 | ΔADH5 | double |
| relative fitness $W$ | 1.000 | 1.000 | 0.999 | 0.699 |
| glucose uptake | 5.00 | 5.00 | 5.00 | 1.98 |
| oxygen uptake | 0.80 | 0.80 | 0.80 | 1.74 |
| ethanol excretion | 8.05 | 8.05 | 8.05 | 0.39 |
| protein, complex III | 1.6 | 1.6 | 1.6 | 3.8 |
| *PRO2* × *QCR2 at* $P_{tot} = 0.19$; $\varepsilon_{ij} = +0.09$ | | | | |
| | wild type | ΔPRO2 | ΔQCR2 | double |
| relative fitness $W$ | 1.000 | 0.871 | 0.658 | 0.658 |
| glucose uptake | 5.00 | 5.00 | 5.00 | 5.00 |
| oxygen uptake | 1.32 | 0.83 | 0.27 | 0.27 |
| ethanol excretion | 7.59 | 8.01 | 8.61 | 8.61 |
| protein, complex III | 2.7 | 1.6 | 0 | 0 |
| protein, CAR2 | 0 | 8.3 | 6.3 | 6.3 |

**Table D. The strict pairs against the genome-wide interaction screen.** Model, $\varepsilon_{ij}$ at the $P_{tot}$ where it is largest (ecYeastGEM-A); screen, the strongest measurement of that gene pair in the non-essential by non-essential array of Costanzo and co-workers [2], with its $p$-value. Scored applies that study's intermediate-confidence criterion, $|\varepsilon| > 0.08$ and $p < 0.05$. The screen is batch culture on glucose, in which the model places the cell in the respirofermentative regime. GPM1, whose pairs carry the sign inversion, is absent from the screen in every allele and so does not appear.

| pair | model $\varepsilon_{ij}$ | screen $\varepsilon$ | $p$ | scored |
|---|---|---|---|---|
| ADH5 × ADH1 | −0.428 | −0.0934 | $1 \times 10^{-5}$ | negative |
| SDH4 × YJL045W | −0.082 | +0.0365 | 0.309 | not scored |
| YJL045W × SDH1 | −0.082 | +0.0796 | 0.033 | not scored |
| SDH4 × SHH4 | −0.082 | −0.0139 | 0.423 | not scored |
| NDE2 × NDE1 | −0.046 | +0.0449 | 0.004 | not scored |
| GND2 × GND1 | −0.037 | — | — | not measured |
| SOL4 × SOL3 | −0.037 | −0.1331 | $1 \times 10^{-57}$ | negative |
| ALT2 × ARO8 | −0.032 | +0.0125 | 0.410 | not scored |
| ILV1 × JEN1 | −0.021 | −0.0099 | 0.334 | not scored |
| GNP1 × JEN1 | −0.021 | −0.0251 | 0.153 | not scored |
| YAT1 × YAT2 | −0.019 | −0.0105 | 0.403 | not scored |
| ALT2 × ARO9 | −0.019 | −0.0139 | 0.344 | not scored |
| PET9 × AAC1 | −0.018 | — | — | not measured |

### References

[1] van Hoek P, van Dijken JP, Pronk JT. Effect of specific growth rate on fermentative capacity of baker's yeast. Appl Environ Microbiol. 1998;64(11):4226-33. doi:10.1128/AEM.64.11.4226-4233.1998.

[2] Costanzo M, VanderSluis B, Koch EN, Baryshnikova A, Pons C, Tan G, et al. A global genetic interaction network maps a wiring diagram of cellular function. Science. 2016;353(6306):aaf1420. doi:10.1126/science.aaf1420.
